# Evolving the olfactory system with machine learning

**DOI:** 10.1101/2021.04.15.439917

**Authors:** Peter Y. Wang, Yi Sun, Richard Axel, L.F. Abbott, Guangyu Robert Yang

## Abstract

The convergent evolution of the fly and mouse olfactory system led us to ask whether the anatomic connectivity and functional logic of olfactory circuits would evolve in artificial neural networks trained to perform olfactory tasks. Artificial networks trained to classify odor identity recapitulate the connectivity inherent in the olfactory system. Input units are driven by a single receptor type, and units driven by the same receptor converge to form a glomerulus. Glomeruli exhibit sparse, unstructured connectivity to a larger, expansion layer of Kenyon cells. When trained to both classify odor identity and to impart innate valence onto odors, the network develops independent pathways for identity and valence classification. Thus, the defining features of fly and mouse olfactory systems also evolved in artificial neural networks trained to perform olfactory tasks. This implies that convergent evolution reflects an underlying logic rather than shared developmental principles.

## Introduction

The anatomic organization and functional logic of the olfactory systems of flies and mice are remarkably similar despite the 500 million years of evolution separating the two organisms. Flies and mice have evolved odorant receptors from different gene families and employ distinct developmental pathways to construct a similar neural architecture for olfaction, suggesting that the similarity between the two olfactory systems emerged by convergent evolution. The sensory neurons in each organism express only one of multiple odor receptors. This singularity is maintained with the convergence of like neurons to form glomeruli so that mixing of olfactory information occurs only later in the processing pathway. Convergent evolution of the olfactory system may reflect the independent acquisition of an efficient solution to the problems of olfactory perception. We asked whether networks constructed by machine learning to perform olfactory tasks share the organizational principles of biological olfactory systems.

Artificial neural networks (ANNs) (Lecun, Bengio, & Hinton, 2015) capable of performing complex tasks provide a novel approach to modeling neural circuits (Mante, Sussillo, Shenoy, & Newsome, 2013; Yamins & DiCarlo, 2016). Neural activity patterns from higher visual areas of monkeys viewing natural images resemble activity patterns from neural networks trained to classify large numbers of visual images (Yamins & DiCarlo, 2016). These results reveal a correspondence between the artificial and biological visually driven responses. However, it has been difficult to determine to what extent the connectivity of ANNs recapitulates the connectivity of the visual brain. Multiple circuit architectures can be constructed by machine-learning methods to achieve similar task performance, and details of connectivity that might resolve this ambiguity remain unknown for most mammalian neural circuits. In contrast, the precise knowledge of the connectivity of the fly olfactory circuit affords a unique opportunity to determine whether ANNs and biological circuits converge to the same neural architecture for solving olfactory tasks. In essence, we have used machine learning to ‘replay’ evolution, to explore the rationale for the evolutionary convergence of biological olfactory circuits.

In fruit flies, olfactory perception is initiated by the binding of odorants to olfactory receptors on the surface of sensory neurons on the antennae (Figure 1a). Individual olfactory receptor neurons (ORNs) express one of 50 different olfactory receptors (ORs), and all receptor neurons that express the same receptor converge onto an anatomically invariant locus, a glomerulus within the antennal lobe of the fly brain (Vosshall, Amrein, Morozov, Rzhetsky, & Axel, 1999; Vosshall, Wong, & Axel, 2000a). Most projection neurons (PNs) innervate a single glomerulus and send axons to neurons in the lateral horn of the protocerebrum (LHNs) and to Kenyon cells (KCs) in the mushroom body (MB) (Jefferis et al., 2007; Marin, Jefferis, Komiyama, Zhu, & Luo, 2002; Wong, Wang, & Axel, 2002). The invariant circuitry of the lateral horn mediates innate behaviors (Datta et al., 2008; Jefferis et al., 2007; Tanaka, Awasaki, Shimada, & Ito, 2004), whereas the MB translates olfactory sensory information into associative memories and learned behaviors (De Belle & Heisenberg, 1994; Dubnau, Grady, Kitamoto, & Tully, 2001; Heisenberg, Borst, Wagner, & Byers, 1985; McGuire, Le, & Davis, 2001).

**Fig. 1.**
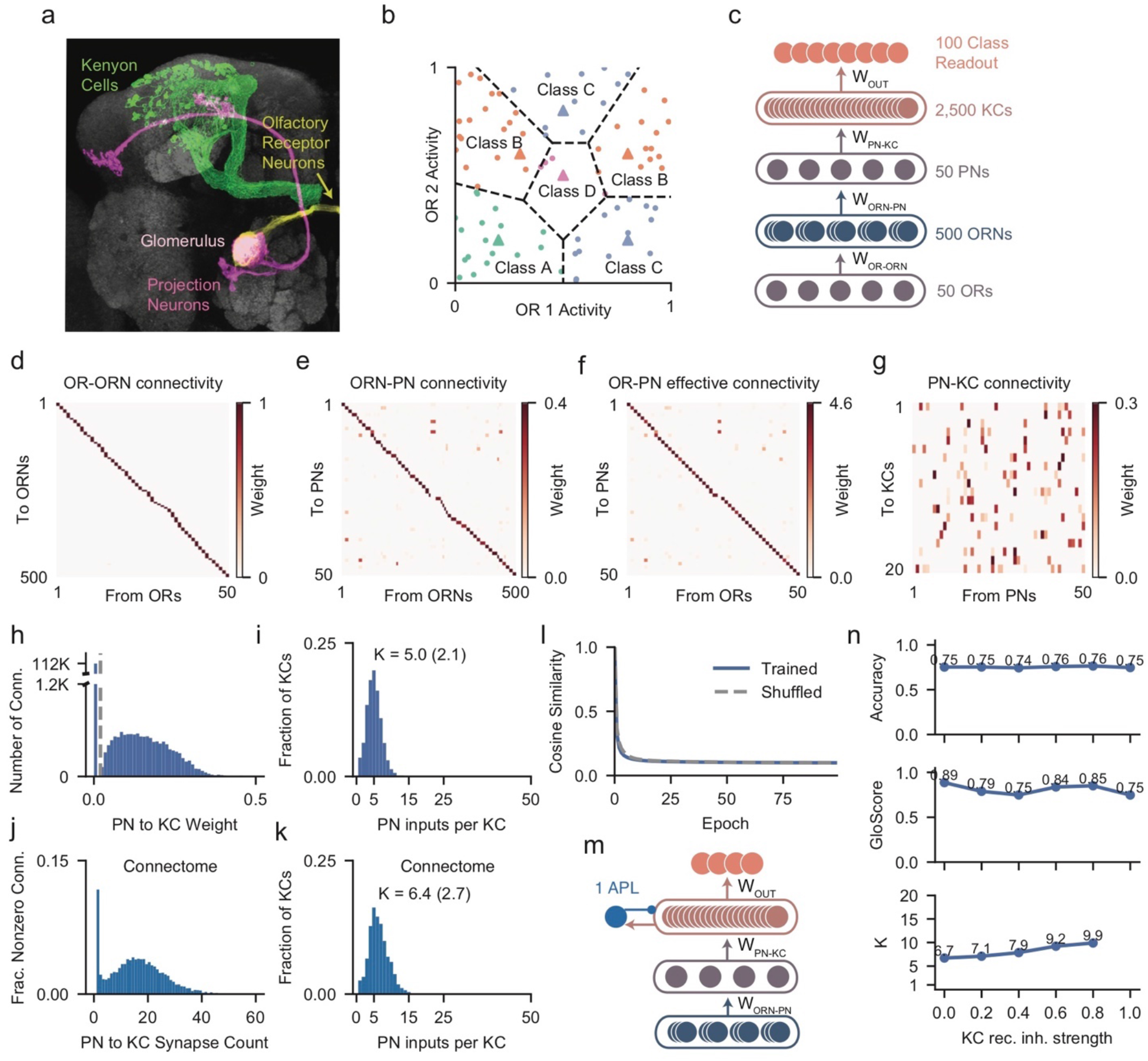
Artificial neural network evolves the connectivity of the fly olfactory system. **a**. The fly olfactory system. **b**. Illustration of the task. Every odor (a million in total, 100 shown) is a point in the space of ORN activity (50 dimensions, 2 dimensions shown) and is classified based on the closest prototype odor (triangles, 100 in total, 4 shown). Each class is defined by two prototype odors. **c**. Architecture of the artificial neural network. The expression profile of ORs in every ORN as well as all other connection weights are trained. **d**. OR-ORN expression profile after training. ORNs are sorted by the strongest projecting OR. **e**. ORN-PN mapping after training. Each PN type is sorted by the strongest projecting ORN. **f**. Effective connectivity from OR to PN type, produced by multiplying the matrices in (**d**) and (**e**). **g**. PN-KC connectivity after training, only showing 20 KCs (2500 total). **h**. Distribution of PN-KC connection weights after training showing the split into strong and weak groups. Connections weaker than a set threshold (dotted gray line) are pruned to zero (left peak). **i**. Distribution of KC input degree after training. Text near peak shows mean and standard deviation. *K* is the average number of PN inputs per KC. **j**, Distribution of PN-KC synapse counts from the fly hemibrain connectome (Li et al., 2020). **k**, Distribution of KC input degree from the connectome data. Left peak corresponds to connections with one synapse. **l**. Average cosine similarity between the weights of all pairs of KCs during training. At every epoch, the cosine similarity was also computed after shuffling the PN-KC connectivity matrix. This shuffling preserves the number of connections each KC receives but eliminates any potential structured PN inputs onto individual KCs. **m,n**. Investigating the impact of a recurrent inhibitory neuron in the KC layer. **m**, Schematics of a network with a recurrent inhibitory neuron at the KC layer, modeling the Anterior Paired Lateral (APL) neuron. The recurrent inhibitory neuron receives uniform excitation from all KC neurons, and inhibits all KC neurons uniformly in return. **n**. (Top to bottom) Accuracy, GloScore, and KC input degree (*K*) for networks with different strengths of KC recurrent inhibition. Stronger KC recurrent inhibition moderately increases KC input degree while having no clear impact on accuracy and GloScore. *K* value is not shown for the network where KC input degree cannot be reliably inferred (Methods).

Individual Kenyon cells, the intrinsic neurons of the MB, receive unstructured input from ~4-10 PNs (Caron, Ruta, Abbott, & Axel, 2013; Li et al., 2020; Zheng et al., 2018) and densely innervate MBONs, the extrinsic output neurons of the mushroom body (Aso et al., 2014b; Caron et al., 2013; Chia & Scott, 2019; Hattori et al., 2017; Li et al., 2020; Tanaka et al., 2004; Zheng et al., 2018). Synaptic plasticity at the KC-MBON synapse results in olfactory conditioning and mediates learned behaviors (Cohn, Morantte, & Ruta, 2015; Felsenberg et al., 2018; Handler et al., 2019; Hige, Aso, Rubin, & Turner, 2015).

The anatomic organization and functional logic of the mouse olfactory system is remarkably similar to the fly olfactory circuit. Sensory neurons in the mouse express only 1 of ~1000 odorant receptors (Buck & Axel, 1991; Godfrey, Malnic, & Buck, 2004; Zhang & Firestein, 2002). Neurons expressing a given receptor converge onto topographically fixed glomeruli in the olfactory bulb, the vertebrate equivalent of the antennal lobe (Mombaerts et al., 1996; Ressler, Sullivan, & Buck, 1993, 1994; Vassar et al., 1994). The mouse projection neurons, mitral and tufted cells, project to primary olfactory cortex where they synapse onto ~1 million piriform neurons (Price & Powell, 1970). Piriform neurons receive roughly 30-100 inputs from an apparently random collection of glomeruli (Davison & Ehlers, 2011; Miyamichi et al., 2011). The hemi-brain connectome of the fly brain (Scheffer et al., 2020) reports numerous axonal-axonal synapses between Kenyon cells in the mushroom body, but these are not believed to be functional (Li et al., 2020). In contrast, pyramidal cells of the piriform cortex make functional recurrent connections with other excitatory neurons (Franks et al., 2011). These recurrent connections are important for concentration-invariant odor coding (Bolding & Franks, 2018; Stern, Bolding, Abbott, & Franks, 2018) and may shape odor tuning during passive odor experience and learning (Pashkovski et al., 2020; Schoonover, Ohashi, Axel, & Fink, 2021).

The convergent evolution of the fly and mouse olfactory systems led us to ask whether the anatomic connectivity and functional logic of olfactory circuits would evolve in artificial neural networks constructed to perform olfactory tasks. We used stochastic gradient descent (Bottou, 2010; Kingma & Ba, 2014; Lecun et al., 2015; Rumelhart, Hinton, & Williams, 1986) to construct artificial neural networks that classify odors. In trained networks, we found singularity of receptor expression, convergence to form glomeruli, and divergence to generate sparse unstructured connectivity that recapitulate the circuit organization in flies and mice. We found that a three-layer input-convergence-expansion structure is both necessary and sufficient for the odor classification tasks we have considered. We also trained neural networks to classify both odor class and odor valence. After training, an initially homogeneous population of neurons segregated into two populations with distinct input and output connections, resembling learned and innate pathways. These studies provide a logic for the functional connectivity of the olfactory systems in evolutionarily distant organisms.

## Results

### Artificial neural networks converge to biological structures

We designed a family of odor classification tasks that mimic the ability of animals to distinguish between odor classes and to generalize within classes. In the model, each odor elicits a unique pattern of activation across the ORs. Odors are assigned to 100 classes that are defined by odor prototypes. Specifically, each odor belongs to the class of its nearest prototype, measured by the Euclidean distance between receptor activations (Figure 1b). Using only a single prototype to define each class results in a relatively simple olfactory task that can be solved without using the layers of olfactory processing that we wish to explore (Figure S2a-d). Thus, we consider classes that are defined by multiple prototypes, predominantly using two prototypes per class. This means that an odor class corresponds to an association involving multiple different types of odors. We used a training set of a million randomly sampled odors to construct the networks and assessed generalization performance with a test set of 8192 = 2^13^ additional odors.

We first modeled the olfactory pathway as a feedforward network with layers representing 50 ORs, 500 ORNs, 50 PN types, and 2,500 KCs (Figure 1c, Methods). In the following sections, we will consider more realistic network architectures with local interneurons. The model also included a set of 100 output units that allow us to read out the class assigned by the model to a given odor (instead of directly modeling MBONs). The strengths of model connections between the OR and ORN layers represent the levels of expression of the 50 different receptor types in each ORN. ORN-to-PN and PN-to-KC connections represent excitatory synapses between these cell types and are therefore constrained to be non-negative. We chose to represent the ~150 PNs in the antennal lobe as 50 PN types because the ~3 homotypical ‘sibling’ PNs that converge onto the same glomerulus show almost identical activity patterns (Kazama & Wilson, 2009; Masuda-Nakagawa, Tanaka, & O’Kane, 2005). We hereafter refer to PN types as PNs. Initially, all connections were all-to-all and random (Figure 1c), meaning that every ORN expressed every OR at some level and connected to every PN. Similarly, each PN initially connected to all the KCs. Neural responses were rectified linear functions of the total synaptic input, and batch normalization, a process resembling neuronal response adaptation, was applied to PN activity (Methods). The network was trained by altering its connection weights and bias currents with the goal of minimizing classification loss. This occurs when there is high activity only in the readout unit representing the correct class associated which each odor. This process can be thought of as evolving a circuit architecture *in silico*.

Following training of the network, classification was ~75% accurate (chance is ~1%). The initial random, all-to-all connectivity changed dramatically during the training process. After training, all but one of the OR-to-ORN coupling strengths for each OR are close to zero (Figure 1d). This corresponds to the expression of a single OR in each ORN. Similarly, all but ~10 of the ORN connections to each PN approach zero (Figure 1e) and, for each PN, all of these connections arise from ORNs expressing the same OR type (Figure 1e). This recapitulates the convergence of like ORNs onto a single glomerulus and the innervation of single glomeruli by individual PNs (Mombaerts et al., 1996; Vosshall, Wong, & Axel, 2000b). The extent that PNs receive input from a single OR type was quantified by GloScore, which, for each PN, is the difference in magnitude between the strongest two connections it receives from the OR types divided by their sum (Methods). A GloScore of 1 indicates that each PN receives all its inputs from a single OR type, recapitulating fruit fly connectivity. During training of the network, the GloScore of ORN-PN connectivity started near 0 and quickly approached values close to 1 (Figure S1b). Thus, the model recapitulates both the singularity of OR expression in the ORNs and the existence of glomeruli in which ORNs expressing the same OR converge and connect to a glomerulus innervated by a single PN type.

The model also recapitulated distinctive features of PN-to-KC connectivity. Each KC initially received connections from all 50 PNs but, during training, connections from PNs to KCs became sparser (Figure 1g, S1b). To quantify the number of PN inputs that each KC receives, weak PN-to-KC connections were pruned to zero during training (Figure 1h, S1d-e). Results are insensitive to the precise value of the pruning threshold, and pruning did not reduce classification performance (Figure S1d). Furthermore, we found that the average number of PNs per KCs, *K*, plateaued during training, with a sparse *K*~3-7 PN inputs for each KC (Figure 1i, S1b). This closely matches the value (K ~ 6) derived from the hemibrain connectome of the adult fruit fly (Figure 1j,k, Li et al., 2020). Importantly, this sparse connectivity can also be obtained without pruning (Figure S1d, Methods). In some cases, no distinct gap separated weak from strong synapses, making an estimate of connection sparsity ambiguous; we identify these instances when they occur and exclude them from further analysis (Figure S1c).

The sparsity and lack of structure in the PN-to-KC connections of the model recapitulate the properties of these connections in the fly (Caron et al., 2013; Li et al., 2020; Zheng et al., 2018). The sparse KC input had no discernable structure (Figure 1l; Figure S3); the average correlation between the input connections of all pairs of KCs is similar to the correlations obtained by randomly shuffled connectivity at every training epoch (Figure 1l). Thus, from ORs to KCs, the ANNs we have trained to classify odors exhibit connectivity that mirrors the layered circuitry of the fly olfactory system, with individual ORs expressing only 1 of 50 receptors, similar ORNs converging onto single glomeruli, individual PNs receiving input from only a single glomerulus, and KCs receiving sparse and unstructured connections from PNs (Video S1). These results were invariant to model hyper-parameters such as training rate and input noise (Figure S1). Moreover, they were also independent of non-zero activity correlations between different ORs (Figure S4a, b). Uniglomerular PNs and sparse, random PN-to-KC connectivity are necessary for high accuracy (Figure S4c). Forcing each PN to receive inputs from multiple ORs (Figure S4d) or introducing stereotypy in PN-to-KC connections (Figure S4e) both substantially reduces accuracy. In all subsequent modelling experiments, we did not include the OR-to-ORN connectivity; instead, every ORN was constructed to express a single OR.

KCs in the fly are inhibited largely through feedback from a non-spiking interneuron, APL (Aso et al., 2014a; Lin, Bygrave, De Calignon, Lee, & Miesenböck, 2014; Tanaka, Tanimoto, & Ito, 2008). We modeled the APL assuming that it receives excitatory input from all KCs and iteratively provides subtractive feedback inhibition onto every KC (Figure 1m). Feedback inhibition did not strongly influence the number of PN inputs per KC, the formation of glomeruli, or task performance (Figure 1n).

### Dependence of results on model features

We next investigated how our results depend on key biological features in the models. The most critical element for the results we have reported is the restriction to non-negative OR-ORN, ORN-to-PN and PN-to-KC connections. Convergence of ORNs expressing the same OR onto PNs does not occur if connections are not sign-constrained. Gloscores drop if ORN-PN connections are not sign-constrained, although classification accuracy is maintained (Figure 2a). In this case, PNs receive a dense array of inhibitory and excitatory connections from ORN inputs, with the ORN connection patterns received by PNs largely uncorrelated (Figure S5a-d).

**Fig. 2.**
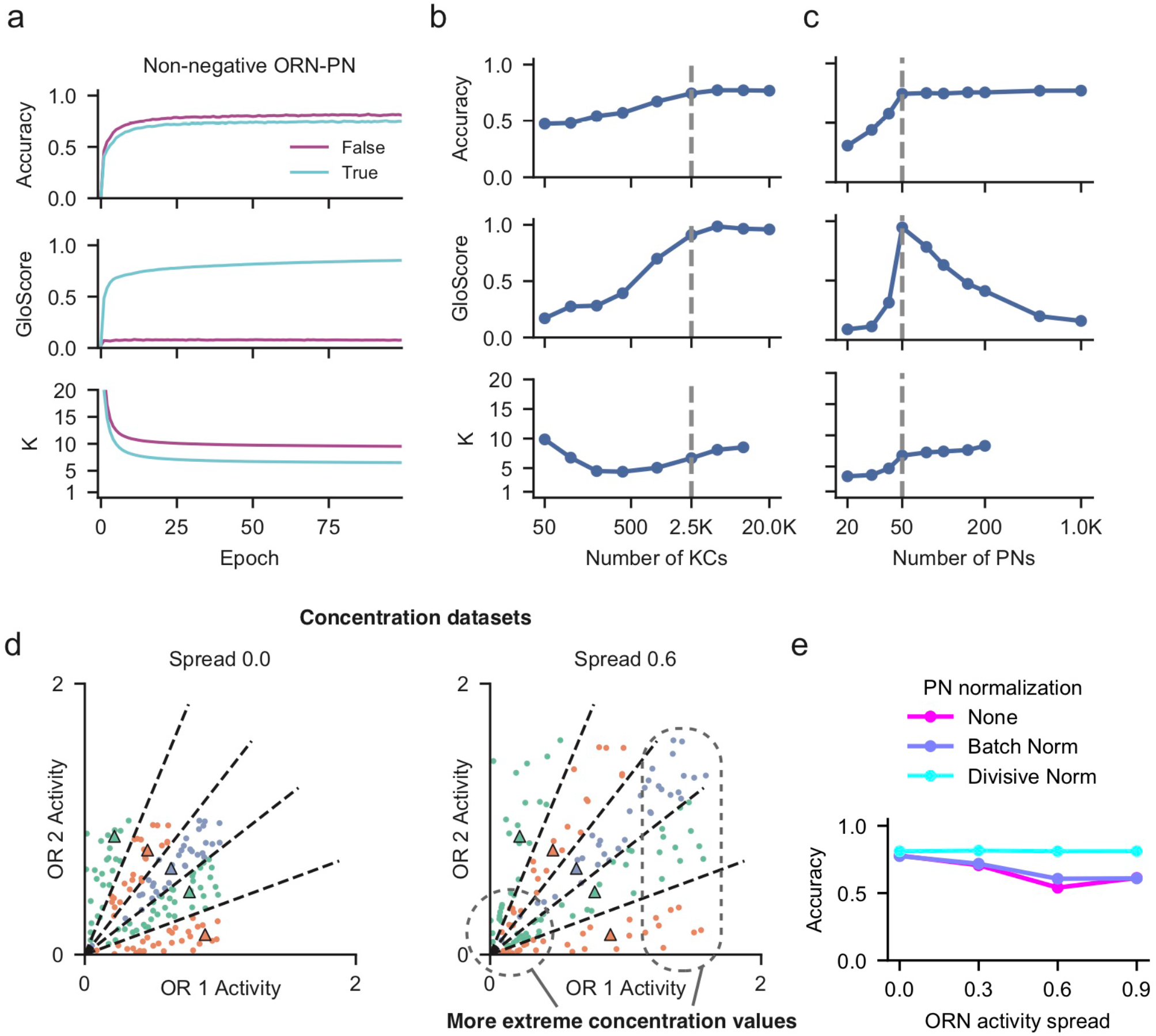
Dependence of results on biological features. (**a**) (From top to bottom) Accuracy, GloScore, and KC input degree as a function of training for networks with and without the non-negativity constraint for ORN-PN connections. **(b,c)** Summary of accuracy, GloScore and KC input degree for trained networks with varying numbers of KCs (**b**), and varying numbers of PNs (**c**). When the number of PNs is high, the KC input degree cannot be reliably inferred. (**d)** Schematics of two concentration-invariant tasks. The odor prototypes (triangles) lie on the unit sphere, making classification boundaries radiate outwards from the origin. The class that each odor belongs to therefore depends on its normalized activity and not on its concentration (i.e., magnitude of OR activity), unlike in the standard task (Figure 1b). (Left) A dataset where each OR’s activity is uniformly distributed across odors. (Right) A dataset where weak and strong odors are more common. The proportion of odors with extreme concentration values is proportional to the “spread”, a parameter between 0 and 1 (see Methods). (**e)** Biological implementations of activity normalization (divisive normalization) rescues classification performance in a concentration-invariant classification task when odor concentration is highly variable. In contrast, a normalization method widely used in machine learning, Batch Normalization (Ioffe & Szegedy, 2015), does not improve performance.

To explore the effect of varying cell numbers, we first trained networks with different numbers of KCs, with ORNs and PNs fixed at 500 and 50, respectively. As the number of KCs was decreased, PNs sampled from multiple ORs, decreasing the GloScore and classification performance (Figure 2b, Figure S5f, S6g-h). Thus, a large expansion layer of KCs is necessary for high classification performance but, with reduced numbers of KCs, some compensatory mixing occurs at the PN level.

We also varied the number of PNs while keeping the numbers of ORNs and KCs fixed at 500 and 2,500, respectively. When the number of PNs is less than the number of unique OR types (50), the PN layer acts as bottleneck and mixing occurs to ensure that all ORs are represented (Figure 2c, Figure S5e, S6a), but performance suffers. When the number of PNs is greater than 50, we observed some PN mixing of ORN input, although this did not improve classification accuracy, which saturates at 50 PNs (Figure 2c, Figure S5e, S6b). A closer examination revealed that PNs segregate into two distinct populations, a population of uni-glomerular PNs receiving a single type of OR and multi-glomerular PNs receiving multiple types of ORs (Figure S6c-f). Moreover, the connection strengths from uni-glomerular PNs to KCs were strong and crucial for classification performance. In contrast, connection strengths from multi-glomerular PNs to KCs were weak, and silencing them minimally impaired classification performance (Figure S6d-e).

Why does a PN layer exist if glomerular connectivity simply copies ORN activity forward to the PNs? Experimental work has shown that the PN layer normalizes odor-evoked responses (Olsen, Bhandawat, & Wilson, 2010), which is likely to be important for classification of odors across a range of concentrations. We trained a feedforward network (Figure 1c) to perform concentration-invariant classification with and without PN normalization while systematically varying the range of odor concentrations in the task dataset (Figure 2d; Methods). We normalized PN activity using a divisive normalization model inspired by the experimental studies (Luo, Axel, & Abbott, 2010; Olsen et al., 2010). As the range of odor concentrations increased, divisive normalization allowed the network to perform concentration-invariant classification (Figure 2e). *K* remains sparse when divisive normalization is introduced (Figure S6i, j), regardless of the range of odor concentrations.

### Recurrent neural networks converge to biological structures

By varying the numbers of PNs and KCs, we found that performance plateaus when the number of PNs (50) matches the number of ORs, and marginal performance gains were observed when the number of KCs was increased past 2500. However, in the models we have considered thus far, the number of neurons in each layer and the number of layers are fixed. We next asked what structure emerges from a neural network that is not only capable of modifying connection strengths, but also capable of allocating the number of neurons per layer.

To remove *a priori* constraints on the numbers of neurons at each layer, we constructed a recurrent neural network model (RNN) in which ‘layers’ are represented by network processing steps (Figure 3a). The RNN receives odor inputs at the first time step and produces classification outputs after several steps of processing. The training algorithm determines how many neurons are active at each processing step, allowing us to infer a particular layered network architecture. This unconventional use of an RNN allowed us to study how finite resources – neurons and their connections – should be distributed across layers, while training only a single network.

**Fig. 3.**
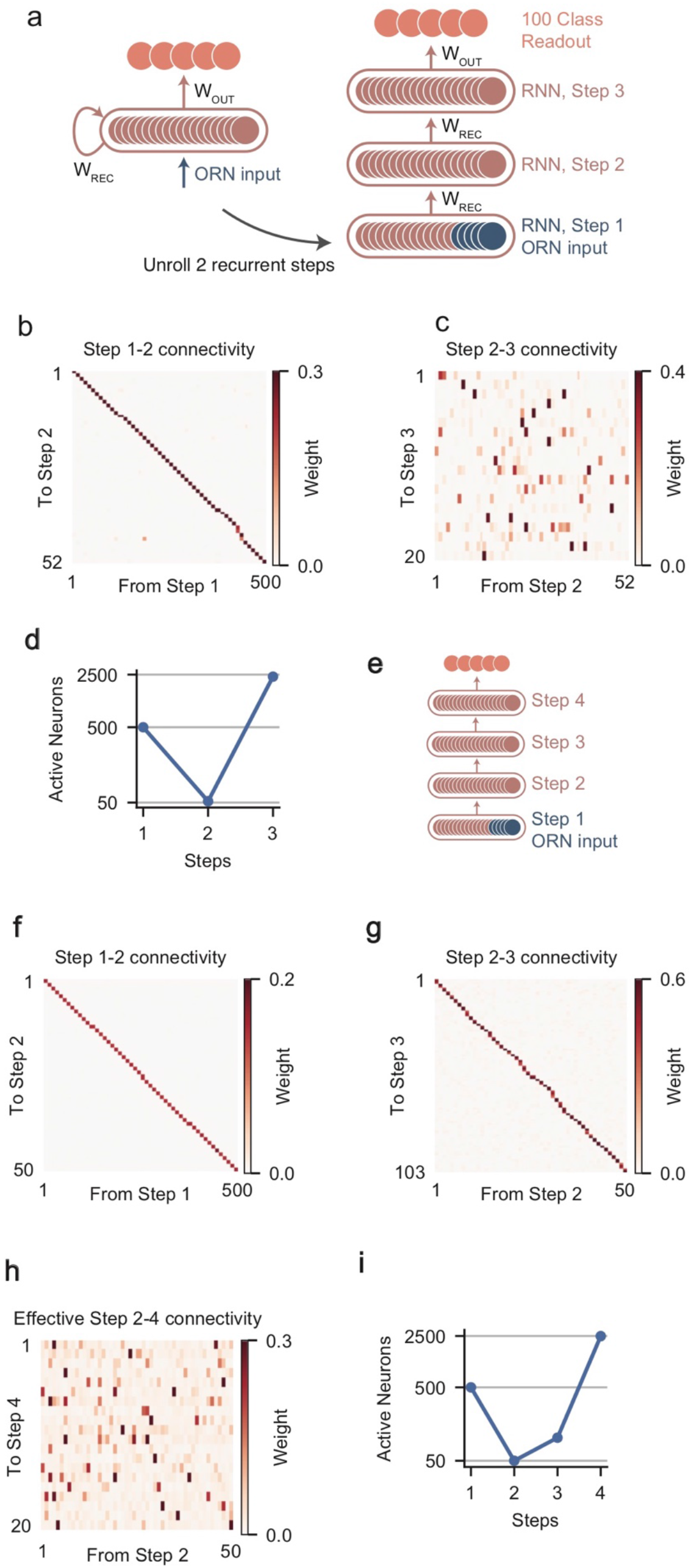
Recurrent neural networks converge to biological structures. **a**. Schematic of a recurrent neural network using recurrent connections (WREC) (left), and the equivalent “unrolled” network diagram (right). **b, c**, Network connectivity between neurons whose activity, when averaged across all odors, exceeds a threshold at different steps. **b**, Connectivity from neurons active at step 1 to neurons active at step 2. Connections are sorted. **c**, Connectivity from neurons active at step 2 to neurons active at step 3, only showing the first 20 active neurons at step 3. **d**. Number of active neurons at each step of computation. At step 1, only the first 500 units in the recurrent network are activated by odors. Classification performance is assessed after step 3. **e-i**, Similar to (**a-d**), but for networks unrolled for 4 steps instead of 3. Classification readout occurs at step 4. Effective step 2-4 connectivity is the matrix product of the step 2-3 (**g**) and step 3-4 connectivity (not shown).

We first considered an RNN in which odor classes were read out after three processing steps (Figure 3a). The RNN model contained 2,500 neurons and was initialized with random, all-to-all, non-negative connectivity between all neurons. At the first processing step, 500 of the 2,500 recurrently connected neurons were provided with OR inputs, and the remainder of the neurons were silent. Thus, this first step of processing represents the ORN layer. After training, this RNN reaches 67% accuracy, slightly lower than that of the feedforward network.

To test whether the RNN self-organized into a compression-expansion structure like the feedforward network, we quantified how many neurons were active at each processing step. Because we did not regularize for activity in the RNN units, a significant number of neurons have non-zero but weak activations to odors (Figure S7a). These levels of activity were bimodally separate from units possessing high levels of activity and were counted as inactive (Figure S7a, Methods). Although the RNN could have used all 2,500 neurons at each processing step, odor-evoked activity from the 500 neurons initialized with ORN activations propagated strongly to only ~50 neurons after the second processing step (Figure 3d). This resulted from the convergence of ORNs onto these PN-like neurons (Figure 3b). In contrast, nearly all neurons of the RNN at the third processing step had average activities (across odors) above the threshold (Figure 3c). These neurons were driven by sparse, unstructured connections from ~5-10 PN-like neurons to the remaining ~2,500 RNN neurons (Figure 3c, Figure S7b-d). Thus, the RNN recapitulated known features of the olfactory circuitry even when the numbers of neurons available at each level was unconstrained.

We next examined the consequence of allowing the RNN to perform four processing steps, which is equivalent to forcing an additional feedforward layer prior to classification of odors (Figure 3e). Interestingly, this network did not use the extra layer to perform additional computations. Rather, it simply copied the activity of the 50-55 PN-like neurons at the second processing step to another similar set of ~100 neurons at the third processing step, only activating the bulk of the 2,500 neurons at the fourth processing step (Figure 3f-i, Figure S7e-h). This result shows that the three-layer olfactory system architecture (input, compression, expansion) is sufficient for the olfactory tasks we considered.

### Network models with ongoing plasticity

We have shown thus far that biological connectivity emerges from both feedforward and recurrent network models when trained on an odor classification task with fixed odor-class mappings. However, the fly olfactory circuit must accommodate the learning of novel odor associations for the fly to adapt successfully to new environments. Evidence strongly suggests that plasticity in synaptic connections from KCs to MBONs underlies olfactory learning (Cohn et al., 2015; Felsenberg et al., 2018; Handler et al., 2019; Hige et al., 2015), whereas PN-KC connection strengths are thought to be fixed (Gruntman & Turner, 2013; Wilson, 2013). We therefore introduced Hebbian plasticity between KCs and class neurons and sought to understand how the KC representation can support ongoing learning. To focus on the PN-KC representation, we eliminated the ORN layer in these studies (Figure 4a).

**Fig. 4.**
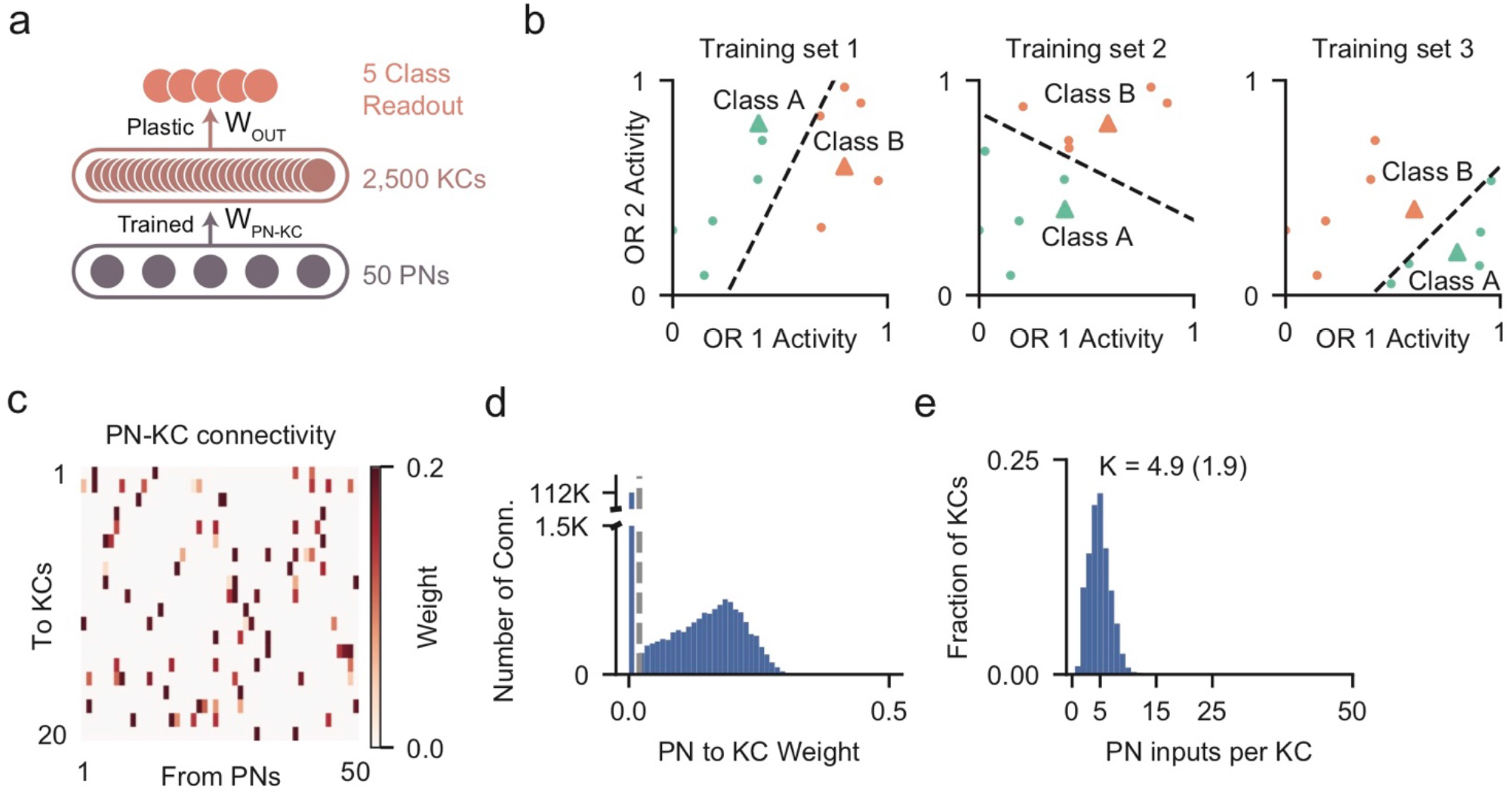
Network models with ongoing plasticity. **a**. Schematic of a meta-trained network. The PN-KC architecture is evolved to support flexible odor learning at the KC-output synapse (W_OUT_). **b**. Multiple datasets are sequentially presented to the network. Each dataset contains a small number of classes and 16 samples from each class. During the presentation of each dataset, KC-output connections undergo rapid plasticity to learn the classes. After fast KC-output learning, generalization performance to a new test set of odors that obey the same classification boundaries is assessed and then used to update, i.e. meta-train, the weights of the network. **c**. PN-KC connectivity after training, showing 20 KCs. **d**. Distribution of PN-KC connection weights after training. **e**. Distribution of KC input degree after training.

Up to this point, networks were trained to assign odors to a fixed set of classes. Now, we construct networks that, after training, can continue to learn new odor classes. This is possible because the networks are expanded to include ongoing plasticity at the synapses between the KCs and output units (Methods). On each episode, we randomly select 16 odors from each of 5 odor classes drawn from the dataset described previously (Figure 1b). During each episode, the feedforward network (Figure 4a) uses synaptic plasticity to learn a new odor-class mapping (Figure 4b) (Finn, Abbeel, & Levine, 2017). After training, the KC-output synapses have undergone plastic updates whereas the remaining network weights are fixed (Methods).

After the update of the plastic synapses, performance for each training episode is assessed by a set of new odors drawn from each one of the 5 odor classes used on that episode, and the non-plastic network weights are adjusted by backpropagation to minimize errors. This encourages the network to generalize to new odors on the basis of a limited set of sampled odors (16-shot learning). At the start of each episode, non-plastic network weights are retained but plastic weights are reset. We asked what connectivity evolved between PNs and KCs to support rapid, flexible learning at the output synapses.

We found that, after training, networks with KC-output plasticity were capable of learning new odor categories. These networks reached up to 80% accuracy in the 16-shot learning task (Figure S8a). Sparse, unstructured connectivity emerged in plastic network models, with an average of ~5 PNs per KC (Figure 4d-e). These results did not depend strongly on hyper-parameters such as the addition of trainable ORN-PN weights, the number of classes per episode, or the number of training odors per class (Figure S8a-c). We conclude that PN-KC connectivity supporting rapid, flexible learning is similar to that observed in the original odor classification task.

### Predicting connection sparsity for different species

The anatomic organization and functional logic of the fly olfactory system is shared with the mouse despite the large evolutionary distance separating the two species. In both mouse and fly, ORNs converge onto a glomerular compression layer, which then projects sparsely to an expansion layer (KCs in the fly, piriform cortex neurons in the mouse). Unlike in the fly, the input degree to the expansion layer in mouse (or any other species) can only been inferred from existing data as *K*~40 − 100 (Davison & Ehlers, 2011; Miyamichi et al., 2011) (Figure 5, Methods).

**Fig. 5.**
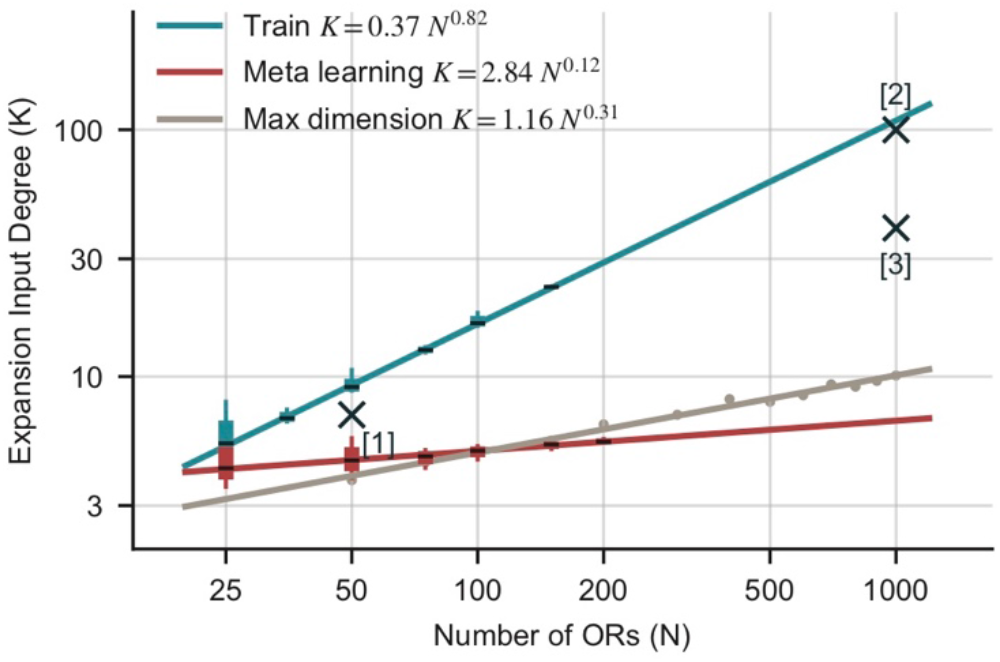
Sparsity for different species. The input degree *K* for networks with different numbers of ORs (*N*). *K* predicted by various methods and is fitted with power-law lines. Cyan: training using the fixed odor categorization task; red: meta-training using the plastic odor categorization task; gray: optimal *K* predicted by maximum dimensionality (Litwin-Kumar et al. 2017); Crosses: Experimental estimates. [2]: Miyamichi et al., 2011; [3]: (Davison & Ehlers, 2011). For each *N*, error bars are derived from networks trained with different learning rates.

We hypothesize that this input degree depends on a variety of parameters, but most heavily on the number of OR types (~1,000 in mouse compared to ~50 in fly). Therefore, in our neural network we asked how the expansion layer input degree (*K*) scales with the number of ORs (*N*), termed *K* − *N* scaling. We have presented networks trained to perform two related yet different tasks, one with a fixed set of odor classes using supervised training and non-plastic synapses (Figure 1), and the other with changing odor classes using meta-training and plastic synapses (Figure 4). Both of these led to similarly sparse PN-KC connectivity in fly-sized networks, *K*~5 − 7 for *N* = 50 (Figure 1i, 4e). We now quantify the *K* − *N* scaling for each of them.

We constructed feedforward network models with different numbers of ORs to examine how their connectivity scales with OR number (Figure S9). Over the range we considered, *K* always increases as a power law function of *N*. However, the *K* − *N* scaling is substantially different across the two tasks. We found that *K* ≈ 0.37*N*^0.82^ for networks trained with fixed classes (Figure 5, blue line), while *K* ≈ 2.84*N*^0.12^ for networks with plasticity (Figure 5, red line). Notably, both scaling results predict qualitatively sparse connectivity since the exponents are significantly lower than 1. The shallower scaling found in plastic networks is broadly consistent with that predicted by previous theoretical work based on determining the wiring that maximizes dimensionality (Figure 5, gray line, Litwin-Kumar et al. 2017). The connectivity that maximizes dimensionality gives rise to *K* ≈ 1.16*N*^0.31^ (Methods).

Although both the fixed and plastic tasks we used to construct networks result in quantitatively similar sparse PN-KC connectivity in fly-sized networks, they make substantially different predictions for mouse-sized networks (*N*~1000): *K* ≈ 0.37 × 1000^0.82^ ≈ 106 for fixed-category training, and *K* ≈ 2.84 × 1000^0.12^ ≈ 7 for the plastic task. Therefore, only fixed-category training appears to produce a result consistent with the mouse data (*K*~40 − 100). However, we note that we have only explored one method to introduce ongoing plasticity. The apparent discrepancy between the mouse data and our plastic network prediction should not be taken as evidence that plasticity and rapid learning of associations are not important in early olfactory processing.

### The emergence of an innate pathway

The repertoire of odorant receptors supports the detection of a vast number of odors in the environment, but a smaller number of receptors exhibit specificity for odors that elicit innate behaviors (Dweck et al., 2015; Ebrahim et al., 2015; Kurtovic, Widmer, & Dickson, 2007; Min, Ai, Shin, & Suh, 2013; Stensmyr et al., 2012; Suh et al., 2004). In flies, PNs activated by these odors project to topographically restricted regions of the lateral horn (LH) to drive innate responses (Datta et al., 2008; Jefferis et al., 2007; Ruta et al., 2010; Varela, Gaspar, Dias, & Vasconcelos, 2019). We asked whether an artificial network could evolve segregated pathways for innate and learned responses.

We trained neural networks to classify both odor class and odor valence. Odor class was determined as in our original models. To add an innate component, each odor was assigned to one of 3 categories, ‘appetitive’, ‘aversive’, or ‘neutral’. Neutral odors activated all ORs as in our previous networks, with activations drawn from a uniform distribution between 0 and 1 (Figure 6a, left). Each odor bearing a non-neutral valence activated all ORs but also a single innate OR especially strongly (on average three times stronger than other ORs). Of the 50 ORs, five were assigned innately appetitive responses, and another five were assigned innately aversive responses. We used a feedforward architecture with 500 ORNs, 50 PNs, and 2,500 third-order neurons that project to both class and innate valence output units (Figure 6b). In this case, there are two sets of output units, one set to report odor class and another to report odor valence. The 2,500 third-order model neurons represent a mixture of LHN and KC neurons, allowing us to investigate whether the segregation into two distinct populations is recapitulated by the model.

**Fig. 6.**
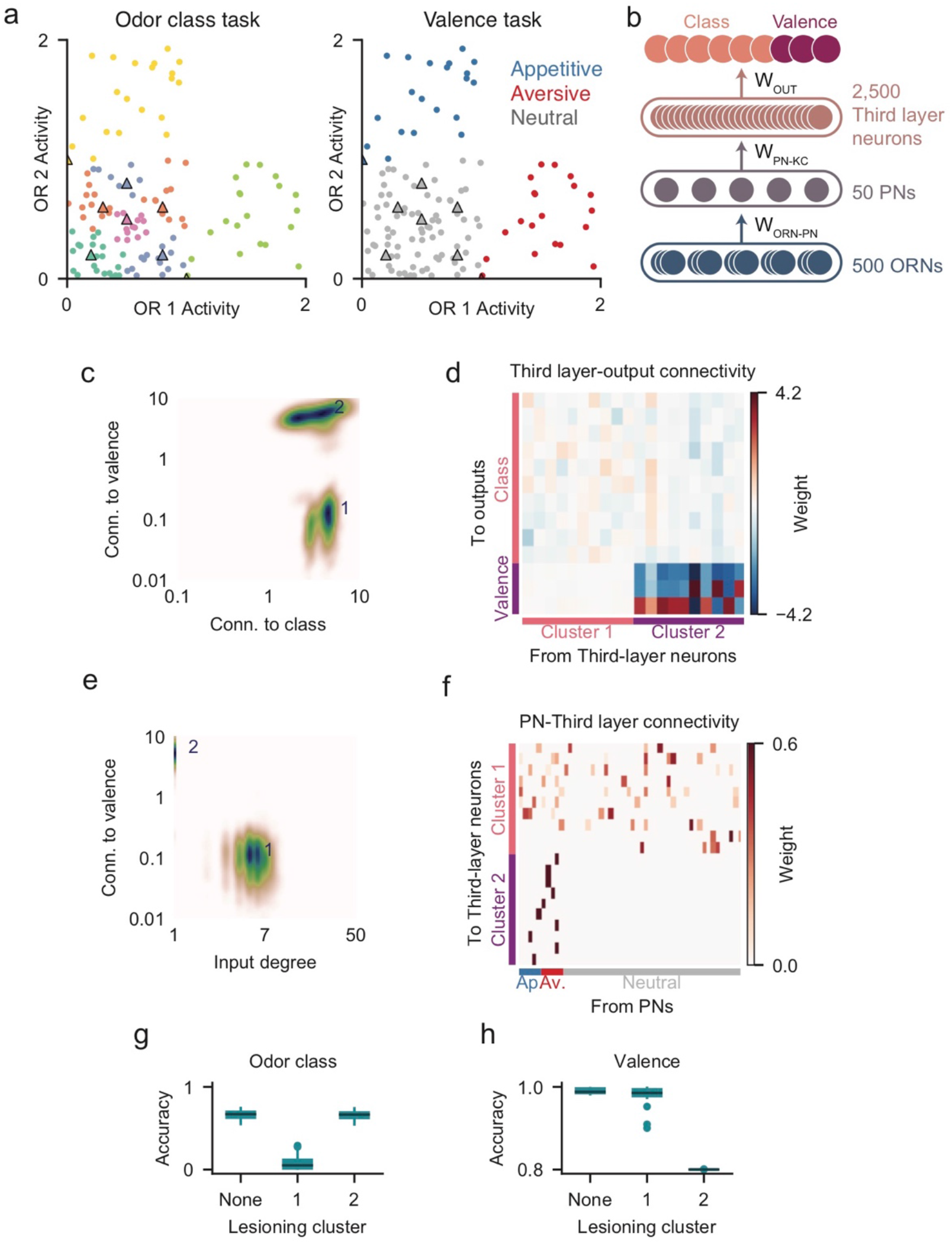
Emergence of separate innate and learned pathways. **a**. Illustration of the class (left) and valence (right) tasks. Non-neutral odors (right, appetitive in blue or aversive in red) each strongly activates one non-neutral OR. The network is trained to identify odor class (left) as previously described (Figure 1b) and also to classify odors into 3 valences (right). **b**, Schematic of a neural network that is trained to identify both odor class and odor valence using separate class and valence read-out weights. **c**. Distribution of third-layer neurons based on output connection strengths to valence read-out neurons against connection strengths to class read-outs. K-means clustering revealed that the third layer can be segregated into two clusters. The density of each cluster is normalized to the same peak value. **d**, The connectivity matrix from the first 10 third-layer neurons from each cluster to output units, the first 10 of 100 class output neurons and all 3 valence output units are shown. **e**, Distribution of third-layer neurons based on output connection strengths to valence read-out neurons against input degree. The distribution of cluster 2 neurons is difficult to see because almost all of them have the same input degree value of 1. **f**, The connectivity matrix from PNs to first 10 third-layer neurons from each cluster. **g, h**. Lesioning the KC-like cluster (group 1) leads to a dramatic drop in odor class performance. (**g**) Lesioning the LH-like cluster (group 2) substantially impaired odor valence performance (**h**).

The network successfully performed both odor classification and valence determination. Glomeruli emerged for neutral, appetitive, and aversive ORs (Figure S10a). The network also generated two segregated clusters of third-order neurons (Figure 6c-d, Figure S10b; Methods). These clusters were segregated based on both input and output connectivity profiles. Cluster 1 typically contains ~2,000 neurons (Figure S10c-d). Cluster 1 neurons are analogous to KCs and project strongly to class read-out neurons but weakly to valence read-out neurons (Figure 6c, d). They receive ~5-7 strong inputs from random subsets of PNs (Figure 6e, f, Figure S10e-f). In contrast, cluster 2 is smaller, containing ~50-200 neurons. Cluster 2 neurons, analogous to LHNs, project strongly to valence read-out neurons (Figure 6c, d), and typically only receive a single strong PN input (Figure 6e, f). Thus, the inputs to the KCs are unstructured whereas the connections to LHN encoding innate valence are valence-specific (Figure 6f). The innate pathway does not emerge if there are no innate odor receptors that respond more strongly to innate odors (Figure S10g-i).

We lesioned each cluster of KC/LHN neurons separately to assess its contribution to odor and valence classification. Lesioning the putative KC cluster (cluster 1) led to a dramatic impairment in odor classification performance (Figure 6g) but left the determination of valence intact (Figure 6h). In contrast, lesioning the putative LH cluster (cluster 2) substantially impaired valence determination (Figure 6h) but had little effect on classification performance (Figure 6g). These results demonstrate that the model network can evolve two segregated pathways analogous to those in the fly.

## Discussion

Network models constructed from machine learning approaches have been used to study the responses of neural circuits and their relationship to circuit function by comparing the activities of network units and recorded neurons (Mante et al., 2013; Masse, Yang, Song, Wang, & Freedman, 2019; Yamins & DiCarlo, 2016; Yamins et al., 2014; Yang, Joglekar, Song, Newsome, & Wang, 2019). Machine learning models generate unit responses and perform the tasks they are trained to do by developing specific patterns of connectivity. It is difficult to perform a detailed comparison of these connectivity patterns with biological connectomes (Cueva, Wang, Chin, & Wei, 2019; Uria et al., 2020) given the limited connectomic data. The current availability of connectome data from flies (Li et al., 2020; Zheng et al., 2018) and the promise of more connectome results in the future make this an opportune time to explore links between biological connectomes and machine learning architectures.

We found that broad network architectures and detailed features of synaptic connectivity shared by the fly and mouse olfactory systems also evolved in artificial neural networks trained to perform olfactory tasks. The observation that machine learning evolves an olfactory system with striking parallels to biological olfactory pathways implies a functional logic to the successful accomplishment of olfactory tasks. Importantly, the artificial network evolves without the biological mechanisms necessary to build these systems *in vivo*. This implies that convergent evolution reflects an underlying logic rather than shared developmental principles. Stochastic gradient descent and mutation and natural selection have evolved a similar solution to olfactory processing.

We constructed feedforward and recurrent networks using stochastic gradient descent. When the feedforward networks were initialized with each ORN expressing all 50 receptors, each ORN evolved to express a single receptor type, recapitulating the expression pattern of ORs in flies and mice. Further, ORNs expressing a given receptor converge on a single PN, and PNs connect with like ORNs to create a glomerular structure. This convergence, observed in both flies and mice, assures that mixing of information across ORs does not occur at early processing layers.

In the network models we studied, each KC initially received input from all 50 PNs but these connections become sparse during training, with each KC ultimately receiving information from ~4-10 PNs, in agreement with the fly circuitry. Although most of our machine modeling was based on the olfactory system in flies, we extrapolated our networks to olfactory systems of far greater size. The results of this extrapolation depended on the task and training procedure. For fixed odor classes, the original task we considered, we obtained an estimate of the number of inputs to piriform neurons from the olfactory bulb, in rough agreement with data from the mouse (40-100).

The architecture of olfactory systems, *in vivo* and *in silico*, is based upon two essential features: converge of a large number of ORNs onto a small number of glomeruli followed by an expansion onto much larger number of third order neurons. Previous theoretical work suggests that a goal of the olfactory system may be to construct a high-dimensional representation in the expansion layer (KCs in the mushroom body or pyramidal cells in the piriform cortex) to support inference about the behavioral relevance of odors (Babadi & Sompolinsky, 2014; Litwin-Kumar et al., 2017). This hypothesis has two important implications for our results.

One results of this previous work is that task performance is proportional to dimensionality when odor classes are learned through synaptic plasticity of a Hebbian form (Litwin-Kumar et al., 2017). In the learning task that we considered, new odor classes were learned through synaptic plasticity that fits into the Hebbian category, so the resulting network should maximize the dimension of the expansion layer odor representation to optimize performance. Indeed, we found that the sparsity of the connections in the resulting networks has a power-law dependence on the number of olfactory receptor types that roughly agrees with the scaling that follows from maximizing dimensionality. However, we obtained a quite different scaling when we trained non-plastic networks on the fixed-class task. Because these networks do not involve Hebbian plasticity, it is not surprising that they exhibit a different degree of sparsity, but we do not currently know of an underlying theoretical principle that can explain the sparsity and scaling we found in the non-plastic case. Interestingly, it is this case that agrees with existing data on the connectivity in the mouse (Davison & Ehlers, 2011; Miyamichi et al., 2011).

Another requirement for achieving maximimum dimensionality is that the representation of odors by the PNs should be uncorrelated (Litwin-Kumar et al., 2017). This provides an explanation for the formation of glomeruli in our network models. The OR activations we used were uncorrelated and, to maximize dimensionality, the transformation from ORs to ORNs and then to PNs must not introduce any correlation. When the weights along this pathway are constrained to be non-negative, the only connectivity pattern that does not induce PN-PN correlations is an identity mapping from OR types to PN output. This is precisely what singular OR expression and OR-specific projection through olfactory glomeruli provides. Interestingly, the results we found suggest that these ubiquitous features of biological olfactory pathways are not simply a consequence of noise robustness, as has been conjectured, but rather arise as the unique solution to eliminating correlations in the glomerular layer to maximize the dimension of the expansion layer.

## Acknowledgements

We thank current and former members of the Axel and Abbott labs, and many colleagues at the Center for Theoretical Neuroscience at Columbia for fruitful discussions. We particularly thank Ashok Litwin-Kumar for discussions and for his guidance. We thank Li Ji-An and Sun Minni for discussions that led to improved training of weight-constrained networks. We thank Jochen Weber, Alexander Antoniades, and John Pellman for building and maintaining the Columbia Axon GPU cluster. This work was supported by the Columbia Neurobiology and Behavior Program (P.Y.W.), Simons Foundation (L.F.A, R.A.), a Simons Foundation junior fellowship (G.R.Y.), NSF NeuroNex Award DBI-1707398 and the Gatsby Charitable Foundation (L.F.A, G.R.Y., P.Y.W.), and the Howard Hughes Medical Institute (R.A.). R.A. is an HHMI investigator.

## Author Contributions

This work is the result of a close collaboration between P.Y.W. and G.R.Y. P.Y.W. and G.R.Y. performed the research. G.R.Y., P.Y.W., and Y.S. performed the analysis of the dependence of K on N (Figure 5). P.Y.W, R.A., L.F.A. and G.R.Y. designed the study, interpreted the results, and wrote the manuscript.

## METHODS

### RESOURCE AVAILABILITY

#### Lead Contact

Further information and requests for resources should be directed to and will be fulfilled by the Lead Contact, Guangyu Robert Yang.

#### Materials Availability

This study did not generate new, unique reagents.

#### Data and Code Availability

- This paper analyzes existing, publicly available data. These datasets are listed in the key resources table. All new data reported in this paper will be shared by the lead contact upon request.
- All original code has been deposited at https://github.com/gyyang/olfaction_evolution and is publicly available. Besides code to reproduce every panel in the paper, we also make available a self-contained Jupyter notebook that reproduces key results and allows easier exploration.
- Any additional information required to reanalyze the data reported in this paper is available from the lead contact upon request.

### METHOD DETAILS

#### Datasets

To generate the standard dataset, we first generated *N*_proto_ = 200 odor prototypes. Each prototype 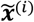 activates *N*_OR_ = 50 ORN types or ORs, and the activation of each ORN type is sampled independently from a uniform distribution between 0 and 1, 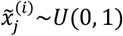. The 200 prototypes are randomly assigned to *N*_class_ = 100 classes, with each class containing two prototypes. A given odor 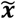 is a vector in the 50-dimensional ORN-type space, sampled the same way as the prototypes. When the network’s input layer corresponds to ORNs, each ORN receives the activation of its OR plus an independent Gaussian noise 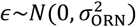, where <_ORN_ = 0 by default (no noise). Its associated ground-truth class *c* is set to be the class of its closest prototype, as measured by Euclidean distance in the ORN-type space. The training set consists of 1 million odors. The validation set consists of 8192 odors.

Besides the standard dataset, we also considered several other datasets based on the standard dataset, as detailed below.

##### Concentration dataset

In this dataset (Figure 2), the prototypes 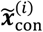 are the normalized versions of the prototypes in the standard dataset 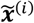, so 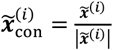. The concentration of each odor is explicitly varied while the average ORN activation across all odors is preserved. For each odor, the activation of each ORN type is sampled from a uniform distribution as described above, and is then multiplied by a concentration scale factor. This scale factor, *s*, is determined by a single parameter, *ϵ*, in which:

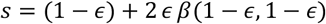

Where *β* is the beta distribution. A value of *ϵ* = 0 produces a dataset with no additional spread, whereas *ϵ* = 1 produces a dataset exhibiting maximal spread with scale factors densely clustered around 0 and 2.

##### Relabel datasets

For the family of relabel datasets (Figure S2), we vary the number of prototypes *N*_proto_=100, 200, 500, 1000 while keeping the number of classes *N*_class_= 100 fixed. We refer to these datasets as relabel datasets, because *N*_proto_ prototypes are relabeled to *N*_class_ classes. The standard dataset uses relabeling as well. The ratio between *N*_proto_ and *N*_class_ is the odor prototypes per class.

##### Meta-learning dataset

This dataset is organized into episodes. Each episode includes a small amount of training data and validation data. In each episode, we randomly select *N*_eps,class_ = 2 classes from the original *N*_class_ 100 classes in the standard dataset. For each of the *N*_eps,class_ classes chosen, we randomly select *N*_eps,sample_ = 16 odors for training and validation respectively. Importantly, within each episode, we re-map each of the *N*_eps,class_ = 2 selected classes to *N*_meta_ = 2 output classes. Intuitively, the network is always doing a (*N*_meta_ =) 2-way classification task. However, the classification boundaries associated with each output class is different in every episode. There is no fixed relationship between the original class label and the new label in each episode, so the network has to learn the new class labels based on the *N*_eps,sample_ data points per class. In total, for each episode, there are *N*_eps,sample_*N*_eps,class_ data points in the training set, and the same amount in the validation set.

##### Valence dataset

In the valence dataset, we replaced *N*_special_ = 10 prototypes from the original *N*_proto_ prototypes with special prototypes that each lies along one axis in the ORN-type space. In other words, each special prototype strongly activates a single ORN type (a special OR), at activity level of 1.0. Of the *N*_special_ special prototypes, *N*_special_/2 = 5 are set to be appetitive or “good” odors, and the other 5 to be aversive or “bad” odors. The rest of the *N*_proto_ − *N*_special_ prototypes and associated odors are set to be neutral and are sampled the same way as the standard dataset. The task is both to classify the odors, as in the standard dataset, and to classify the valence (appetitive, aversive, neutral). In both the training and the validation dataset, we have 10% of the overall odors be appetitive, another 10% be aversive, and the rest 80% be neutral. Therefore, if a network classifies all odors to neutral, the chance level performance for valence classification is 80%. The neutral odors are sampled in the same way as the standard dataset. Each appetitive or aversive odor is sampled by adding the activity level of one special prototype (1.0 for the special OR and 0.0 otherwise) with an activity pattern sampled from a uniform distribution between 0 and 1. In other words, the activity level of an appetitive or aversive odor is sampled randomly from *U*(1, 2) for the special OR, and from *U*(0,1) for other ORs.

##### Correlated dataset

In Figure S4, we introduce correlation between responses of different ORN types. The correlation is independently controlled between 0 and 0.9, while maintaining the marginal distribution of each ORN type to be uniform between 0 and 1. We used a previously proposed method (Cario & Nelson, 1997) for generating such correlated random variables while maintaining their marginal distributions.

#### Network architecture

We train networks of various architectures. The **ORN-PN-KC network architecture** consists of an input layer of 500 model ORNs, 50 PNs, 2500 KCs, and 100 output units. The 500 ORNs are made of 10 ORNs per type for all 50 types of ORNs. The activation of each ORN is the sum of the activation of the corresponding ORN-type 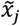 and an independent noise 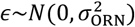, where *σ*_ORN_ = 0 by default (no noise). The ORN-PN, PN-KC, and KC-output connections are all fully-connected at initialization. The ORN-PN and PN-KC connectivity are initialized with a uniform distribution of between 1/*N* and 4/*N*, where *N* is the number of input neurons (500 for ORN-PN, and 50 for PN-KC). The KC-output connectivity is initialized with the standard Glorot uniform initialization. The ORN-PN and PN-KC connections are constrained to be non-negative using an absolute function. All neurons use a rectified-linear activation function (ReLU).

In the **OR-ORN-PN-KC network**, we add an additional layer of OR-ORN connections. Here, the inputs are 50 ORs, activated similarly to the ORNs from the ORN-PN-KC network. The OR-ORN connections are non-negative as well and initialized similarly to ORN-PN and PN-KC connectivity.

For the **identity/valence classification** task, we used a network with two output heads. One containing 100 output neurons as usual. The other contains 3 output neurons for neutral, appetitive, and aversive valence.

We briefly considered an **ORN-Output network** (Figure S2) that has the output directly read out from the ORNs.

Optionally, we include **dropout** on the KC layer, which at training time, but not testing time, set a certain proportion *p*_dropout_ of neuron activities to zero. The default dropout rate is *p*_dropout_ = 0 (no dropout).

The **recurrent network** used in Figure 3 is a discrete-time vanilla recurrent network,

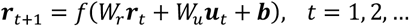

The network consists of 2,500 units. The recurrent connection is initialized uniformly between 0 and 4./2500, the input connection is initialized using Glorot uniform initialization. The recurrent connection is constrained to be non-negative. Out of 2,500 units, 500 receive odor inputs at *t* = 1 in the same way as the ORNs in the feedforward network. The classification output is read-out with at step *T* with connections that are not sign-constrained. By default, we have *T* = 3, which means the network unrolled in time would have 3 layers (*t* = 1, 2, 3) and an output layer.

The **KC recurrent inhibition** mediated by a single APL neuron (Figure 1) is implemented by an inhibitory neuron interacting with the KCs iteratively. The single inhibitory neuron has a neural response equal to the mean KC activation level at each time step. This neuron then sends subtractive inhibitory inputs to all KCs with a connection weight *γ* (KC recurrent inhibition strength in Figure 1). Therefore, the KCs at each time step *t* are activated as

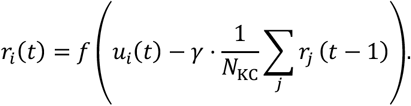

Here *f*(·) is the ReLU activation function. *u_i_*(*t*) = *u_i_* is the feedforward input to the *i*-th unit. *r_i_*(*t*) is the activation level of the *i*-th unit at time step *t*. We run this recurrent inhibition for 10 time steps.

The **divisive normalization** used on the PN layer in Figure 2 is implemented in the following way. Neuron *i* in this layer receives input *u_i_*, and the final activation of this neuron, *r_i_* follows,

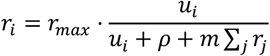

Here, *r_max_*, *ρ*, *m* are parameters that are trained with gradient descent alongside other trainable parameters. In initialization, we have *r_max_* = *N*/2, *ρ* = 0, *m* = 0.99, where *N* is the number of neurons in this layer. For stability during training, we clamped *N*/10 ≤ *r_max_* ≤ *N*, 0 ≤ *ρ* ≤ 3, 0.05 ≤ *m* ≤ 2.

#### Training

The output of the network is linearly read out with trainable weights from the final layer (KC layer in feedforward networks, or the recurrent layer). The loss is softmax cross-entropy loss. The default training method is the adaptive stochastic gradient descent method Adam with learning rate 5e-4, and exponential decay rates for first and second moments 0.9 and 0.999 respectively (the Pytorch default hyperparameter values). The network is typically trained for 100 epochs, each epoch would expose the network to all of the one million odors from the training set.

The training batch size is *B* = 256. By default, we used Batch Normalization on the PN layer to prevent individual neurons from being active or silent for all odors. Technically, Batch Normalization computes the mean *μ_i_* and standard deviation *σ_i_* of inputs *x_i,b_* to the *i*-th single neuron across a minibatch (*b* = 1, …, *B*),

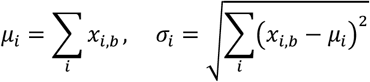

The actual input to the *i*-th neuron is first subtracted by *μ_i_*, then divided by *σ_i_*. It is then multiplied by a trainable parameter, then another trainable parameter is added to it. Biologically, Batch Normalization can be viewed as approximating single neuron adaptation or homeostasis to a range (i.e., a batch) of odors. If a neuron is strongly driven by most odors, then Batch Normalization would reduce its inputs, making this neuron activated in a more balanced manner.

#### Ongoing Plasticity

For the ongoing plasticity results in Figure 4, we use the delta rule to simulate ongoing plasticity in the readout connections (KC-output weights for the model fly network) (Dayan & Abbot, 2005). The delta rule is more biologically plausible than the general gradient descent algorithm because it relies on local information. However, it is not intended to model with high fidelity the biological plasticity rules at the KC-MBON synapses. The delta rule is used here to encourage a KC representation that supports rapid, flexible learning. The default delta rule learning rate is 5e-4.

During each learning episode (see Meta-learning dataset section), the network is presented with a small amount of training and validation data from the meta-learning dataset. The network takes a single delta rule step based on the training data, and the loss is evaluated based on the validation data. The objective of meta-training is to minimize the expected validation loss of the inner training. Meta-training updates all weights and biases in the network at the end of each learning episode using the gradient descent variant, Adam. This meta-training method is a special case of a more general method called MAML, or Model-Agnostic Meta-Learning (Finn, Abbeel, & Levine, 2017a). This method aims at finding (meta-training) parameter values (connection weights and biases) that allow rapid few-step gradient descent learning using a small amount of new training data. We largely adhered to the method detailed in Finn et al., with a few notable exceptions. First, the inner training only performs gradient descent on the KC-output connection. Gradient descent applied only to the last layer reduces to the delta rule. Second, the learning rate of the inner training is allowed to be adjusted by the meta-training process. The latter assumption does not substantially impact our results.

#### Weight pruning and connection sparsity estimation

By default, we have synaptic weight pruning during training. Weights below a certain threshold *θ* are permanently set to zero during and after training. The threshold is set to be *θ* = 1/*N*, where *N* is the number of input neurons for each connectivity matrix. Weight pruning provides a less ambiguous quantitative estimate of connection sparsity.

We observe that in some networks, the distribution of weights has a clear, single peak away from the pruning threshold, and the weight distribution approaches 0 towards the threshold (see Figure S1c for examples). In these cases, the connection sparsity (or density) can be easily inferred by simply quantifying the proportion of connection weights above threshold. However, we found that in some networks (some hyperparameter settings), the distribution of weights has a peak very close to the threshold, making it difficult to count the above-threshold weights. Therefore, we employ a simple heuristic to check if there is a clear peak in the weight distribution far from the pruning threshold. Our heuristic requires the peak of the above-threshold weight distribution be at least 2./*N* larger than the threshold itself, which by default is at 1./*N*. Networks that do not satisfy this “clear peak” criteria are not used to compute the input degree, and their *K* values not shown in plots (e.g. Figure S1a).

When the network does not undergo pruning of weak weights as in some control experiments and for the RNN results, it is necessary to try inferring a threshold separating weak and strong weights. We fit a mixture of two Gaussians model to the log-distribution of weights. The weak/strong weight threshold is where the probability density of the two Gaussian modes cross. In this case, the inferred threshold is used, instead of the pruning threshold, in the above heuristics for determining whether the strong weights have a clear peak in its distribution.

We have done extensive comparisons between networks with and without pruning across various hyperparameter values (many results not shown in figures). For the feedforward network architectures, pruning almost always leads to clearer above-threshold peak in the weight distribution. Importantly, the sparsity result is not a result of pruning per se. When there is no pruning, and the weights clearly separate into weak and strong peaks (for example when *N_proto_* = *N_class_* = 100), the inferred connection sparsity is quantitatively very close to that obtained from networks with pruning. In addition, the network performance is generally identical with or without pruning.

### QUANTIFICATION AND STATISTICAL ANALYSIS

#### GloScore

The glomeruli score (GloScore) of a PN-ORN connectivity matrix *W*_PN→ORN_ is computed by first averaging all connections from ORNs of the same type. For each PN, we find the strongest connection weight *w_1_* and the second strongest connection weight *w_2_* from each ORN type by averaging weights across ORNs of the same type. For non-sign-constrained weights, we use the absolute values of weights. Then GloScore for each unit is computed as,

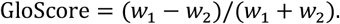

Final GloScore of the entire connection matrix is the average GloScore of all PNs.

#### Inferring connection sparsity from experimental data in mouse

Two previous publications used different approaches to estimate the input degree, K, in mice. The first experiment (Miyamichi et al., 2011) used retrograde anatomic tracing to derive a convergence index of the number of mitral/tufted cells (equivalent of PNs) over the number of piriform neurons (equivalent of KCs), and found values ranging from 3-20. The transfection efficiency of retrograde labeling was estimated to be roughly 10% (Reardon et al., 2016), so the input degree may vary from 30-200 M/T inputs per piriform neuron. The second experiment (Davison & Ehlers, 2011) used optical glutamate uncaging to activate defined points on the olfactory bulb while recording piriform responses, and found that most cells responded to >15 uncaging sites. The authors estimate that 2-3 glomeruli are activated per uncaging site, providing a lower bound of K=40 for input degree.

#### Analysis of Synaptic Connectivity Data from the Hemibrain Connectome

A compact connection matrix summary (v1.2 release) was downloaded from https://www.janelia.org/project-team/flyem/hemibrain. ORNs, uniglomerular, biglomerular and multiglomerular PNs, and KCs and LH neurons were queried according to the naming convention defined in Scheffer et al. 2020. Thermosensory, hygrosensory, and subesophageal zone PNs (VP and Z) were discarded. Given that stronger synapses are formed by increasing the number of synapses, not by larger synapses, as in vertebrates, we use synapse count as a proxy for synaptic strength (Scheffer et al., 2020). Only 2 types of ORNs were present within the dataset, so ORN to PN connectivity was discarded. The distributions of KC input degree and PN to KC synaptic weights were previously reported (Li et al., 2020) and were also extracted from the connectivity of uniglomerular PNs onto KCs. Multiglomerular PNs were excluded because KCs only sample from 0.147 multiglomerular PNs on average.

#### Randomness

To determine whether the frequency of PN input onto KCs is significantly above or below chance expectations, PN-KC connections in the trained network were shuffled while maintaining the number of connections each KC receives. We generated the shuffled data by making a list of PNs that contributed to each PN-KC connection. We then randomly permuted this list and drew from it sequentially to construct a new set of connections for each of the 2500 KCs, drawing as many random connections for each KC as it receives in the trained network. This shuffling eliminates any potential, non-random PN inputs onto individual KCs, and is used to analyze whether KCs are connected to any preferential pair of glomeruli (Figure S3).

To determine whether the distribution of PN inputs onto KCs is binomial, the probability of a connection between each PN with each KC is sampled independently from a Bernoulli distribution with the overall PN-KC connection probability, *p*, of a trained network.

#### Analysis of RNNs

In Figure 3, we analyzed a recurrent neural network, which unlike traditional recurrent networks, is not running in time. Instead we use it as a way to force a limited budget on the total number of neurons, without specifying the exact number of neurons to be used at each processing step.

The key analysis is to infer how many neurons are assigned by the network to each processing step (the same neuron may be used at multiple steps). For each neuron, we computed its average activity at each processing step in response to all odors shown the network. If its average activity at a processing step exceeds a certain threshold, we deem this neuron active at this step. Note that by this definition, an “active neuron” may not be active for each odor. All we ask is that it is sufficiently active for some odors. We used the same threshold of 0.2 across all processing steps, manually chosen after inspecting the distribution of activity (Figure S7). We did not use a threshold of 0 because many neurons are activated very weakly but above zero on average. With positive connection weights and no regularization, it is generally more difficult to have a neuron be activated at 0 across all odors at a given processing step.

#### Analyzing networks of different numbers of OR types

For Figure 5, we trained networks with different numbers of OR types (*N*), ranging from 25 to 200. For simplicity, we focused on the connections from the compression to the expansion layer, while ignoring the connections from ORNs to the compression layer. Therefore, all networks consist of *N* input neurons representing ORN activity, which in turn project to *M* expansion layer neurons. For each value of *N*, the number of expansion layer numbers *M* is set as *N*^2^. For each number of OR, we trained networks with different levels of learning rate 1e-3, 5e-4, 2e-4, 1e-4. We include in our summary plot (Figure 5) only networks that contain a clear peak in the weight distribution, using the criteria established above.

To obtain the maximum dimensionality curve in Figure 5, for each number of OR, we first computed the representation dimensionality (Litwin-Kumar et al. 2017) in response to the training odors when the third-layer input degree is fixed at different values. Then we identified the input degree corresponding to the maximum dimensionality. Finally, we repeat this process for networks with different numbers of ORs. Importantly, we did not use feedforward inhibition that sets the overall mean input to be zero. When mean-canceling feedforward inhibition is used, the maximum dimensionality is achieved at *K* = *N*/2. When introducing an additional constraint on the total number of connections, the optimal *K* becomes substantially lower, around 7 for *N* = 50. However, since we do not constrain the total number of connections for each network, we did not include feedforward inhibition in Figure 5, leading to a *K* that is around 3 for *N* = 50.

#### Analysis of identity/valence two-task networks

For the two-task networks, we used all combinations of the following hyperparameter values: PN normalization (None or Batch Normalization), learning rate (1e-3, 5e-4, 2e-4, 1e-4), KC dropout rate (0, 0.25, 0.5), resulting in 24 networks trained.

To assess whether the expansion layer neurons break into multiple types when analyzing the two-task networks, we represent each third-layer (expansion layer) neuron with three variables: (1) its input degree (the number of above-threshold connections from the previous layer), (2) the norm of its connection weights to the identity classification head, (2) the connection weight norm to the valence classification head. Since these variables are of different scales, we z-scored them (mean subtract then divide by standard deviation). We then obtained a 3-dimensional depiction of each third layer neuron.

Next we did k-means clustering on the normalized data with *k* (the pre-determined number of clusters) ranging from 2 to 10. We quantified the quality of each clustering result with its silhouette score (the higher the better), which intuitively compares the inter-cluster distance with the intra-cluster distance. We found that the optimal number of clusters is generally 2 or 3. We analyzed all networks with 2 optimal clusters. We named the cluster of neurons with stronger connections to the identity readout head as cluster 1, the other as cluster 2.

In Figure 6c, e, we computed the density of neurons in these data spaces separately for each cluster, before adding the densities together. This visualization allows for a clearer depiction of the density peak of each cluster.

When lesioning either cluster 1 or 2 in Figure 6g, h, we set the outbound weights from the lesioned neurons to 0, equivalent to setting their activity to 0.

### KEY RESOURCES TABLE

| REAGENT or RESOURCE | SOURCE | IDENTIFIER |
| --- | --- | --- |
| *Deposited Data* |  |  |
| Hemibrain Connectome compact connection matrix v1.2 | Scheffer et al., 2020 | https://www.janelia.org/project-team/flyem/hemibrain |
| *Software and Algorithms* |  |  |
| Python | Python Software Foundation | https://www.python.org/ |
| Tensorflow | Abadi et al., 2016 | https://www.tensorflow.org/ |
| Pytorch | Paszke et al., 2019 | https://pytorch.org/ |
| SciPy | Virtanen et al., 2020 | https://www.scipy.org/ |
| Numpy | Oliphant, 2006 | https://www.numpy.org/ |
| Scikit-learn | Pedregosa et al., 2011 | https://scikit-learn.org/stable/ |
| Custom code for generating datasets, training networks, analyzing results, and plotting figures | This paper | https://github.com/gyyang/olfaction_evolution |

---

**Figure S1.**
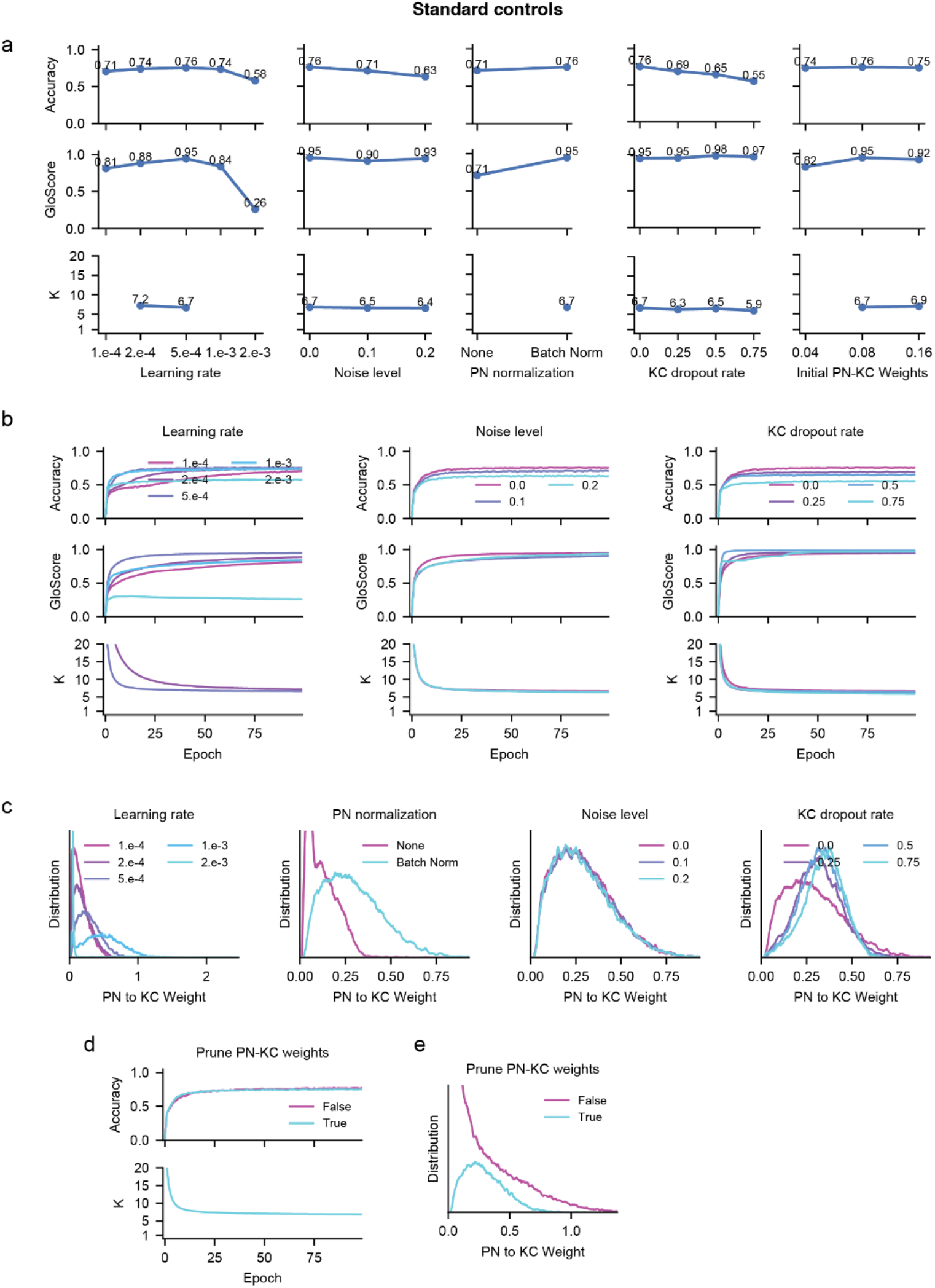
Robust formation of glomeruli and sparse connectivity, Related to Figure 1. **a.** Accuracy (top), GloScore (middle), and KC input degree (K, bottom) as a function of hyperparameters. From left to right, learning rate, noise level, PN normalization, KC dropout rate, and initial PN-KC weights. K values are not shown for networks where the PN-KC connectivity does not contain a single peak well separated from the pruning threshold (see Methods). When a single peak can be inferred (blue dots), GloScore remains high, and KC input degree remains around 5 to 10. **b**. Accuracy (top), GloScore (middle), and KC input degree (K, bottom) during training. For each plot, one hyperparameter is varied, from left or right: learning rate, noise level, and KC dropout rate. Networks of different hyperparameter values converge to the same GloScore and KC input degree during training, as long as PN-KC connectivity is well separated. **c**. Distribution of PN-KC connection weights for networks of different hyperparameter values. Left to right: Learning rate, PN normalization, noise level, and KC dropout rate. Having no PN normalization leads to PN-KC weights poorly separated from the threshold, explaining why in (a) the K value is not shown for the network with no PN normalization. **d, e**. The effect of pruning weak PN-KC weights. Pruning weak PN-KC weights does not affect performance (**d**), but it allows a cleanly separated distribution of PN-KC weights from the threshold (**e**). The lack of a clean separation without pruning (e) leads to unreliable estimation of the PN-KC input degree (d).

**Figure S2.**
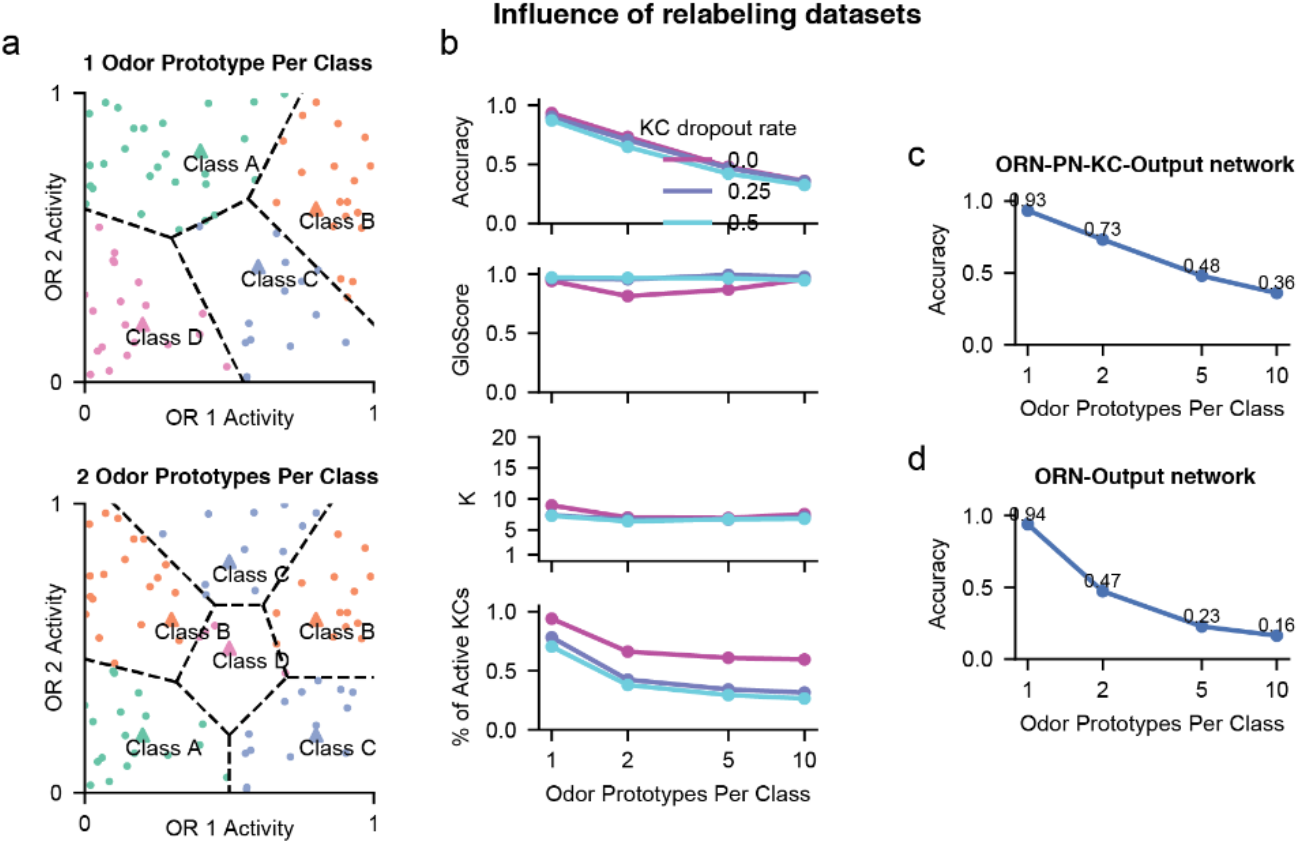
Effect of having multiple odors associated with each class, Related to Figure 1. **a**, Schematics of two datasets. (Top) Illustrating a dataset where only one odor prototype (triangle) is associated with each class. Each class then corresponds to a contiguous area in the input activity space, and the dataset is linearly separable. (Bottom) Illustrating a dataset where each class is associated with two odor prototypes residing in segregated locations in OR activity space. **b**, From top to bottom, accuracy, GloScore, KC input degree, and KC activity sparsity (percentage of KCs active on average) for networks trained on datasets with different numbers of odor prototypes per class. Having more odor prototypes per class promotes KC activity sparsity, while keeping GloScore high and KC input degree almost constant. Having KC dropout has a similar impact. **c, d**, Comparing accuracy between the full ORN-PN-KC-Output network (**c**) and a simple ORN-Output network (**d**). The ORN-Output network has classification readout directly from the ORN input layer. This shallow network performs well on the dataset with 1 odor prototype per class (linearly separable), but much worse than the full network on more complex datasets.

**Figure S3.**
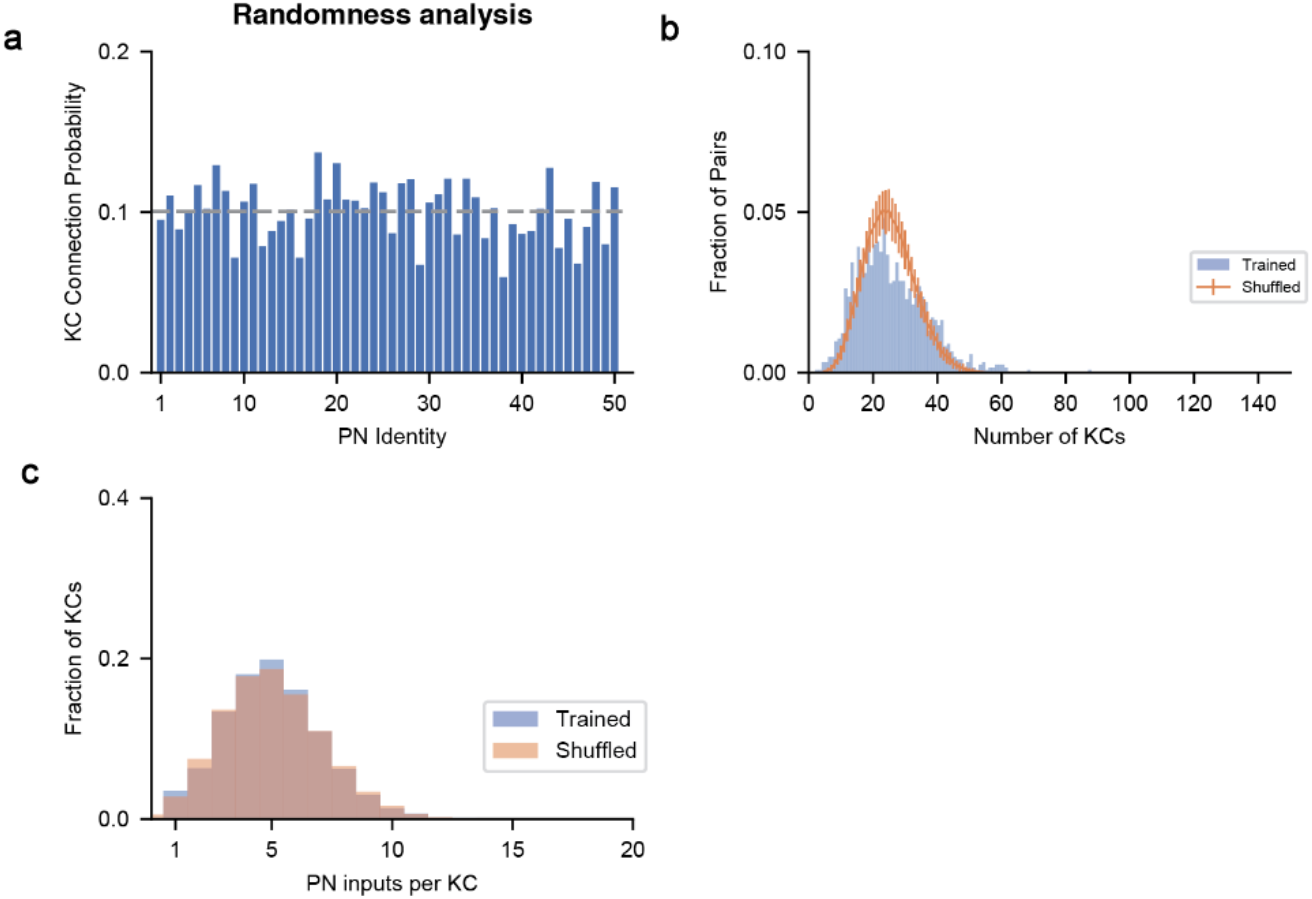
Random sampling of PN inputs from KCs and the impact of KC recurrent inhibition, Related to Figure 1. **a**, Average connection probability from each individual PN (n=50) to all 2500 KCs. KCs sample uniformly from all PNs. **b**, Distribution of number of KCs that receive each of the 1225 (=50×49/2) unique pairs of glomeruli. Data derived from training is shown in blue, and shuffled connections are in orange. Shuffling maintains the frequency of glomerular connections and the distribution of KC input degrees, but eliminates non-random patterns of inputs onto individual KCs. KCs are not preferentially connected to any specific pair of PNs. **c**, Distribution of KC input degrees for all KCs (n=2500). Data derived from training is in blue, and a binomial distribution using the average PN-KC connectivity derived from training data is shown in orange.

**Figure S4.**
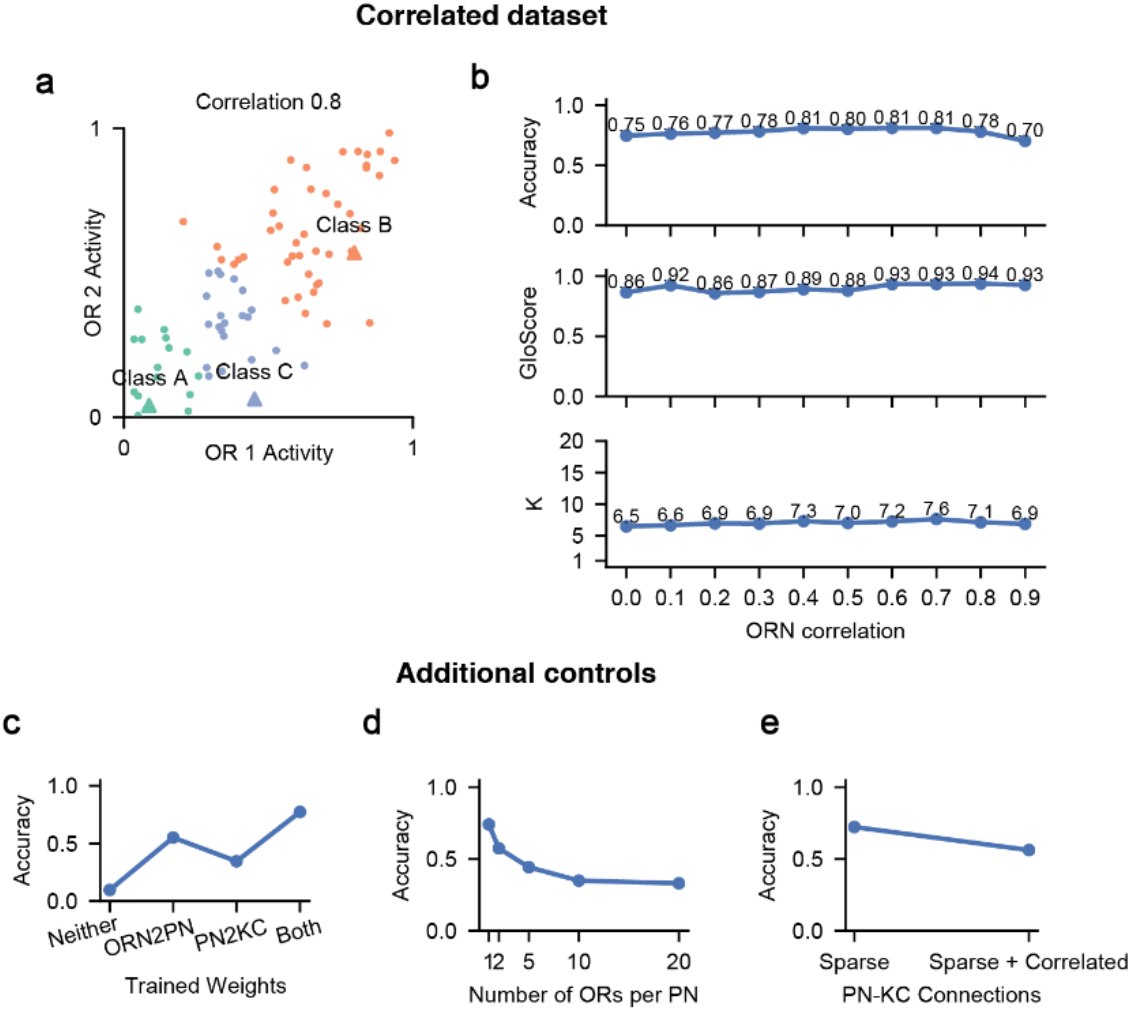
The impact of input correlation and additional controls, Related to Figure 1. **a**, Illustration of a dataset with a 0.8 correlation coefficient between the activities of all OR pairs. **b**, (Top to bottom) Accuracy, GloScore, and KC input degree for networks trained on datasets with different OR correlations. OR correlation level has no clear impact on GloScore and KC input degree. **c**, Task accuracy across several networks in which weights at specified layers are fixed at their random initial values. Fixing connectivity to be random in either one or both layers reduces task performance compared to standard training. In all four scenarios, KC-output weights are trained. Neither: ORN-PN and PN-KC weights are fixed. ORN2PN: ORN-PN weights are trained and PN-KC weights are fixed. PN2KC: ORN-PN weights are fixed and PN-KC weights are trained. Both: ORN-PN and PN-KC weights are trained. **d**, Task accuracy across networks where the number of ORs received by each PN is fixed. In other words, ORN-PN connections are fixed to be multi-glomerular before training. PN-to-KC and KC-output weights are trained, and performance was assessed after training. Classification performance degrades as PNs receive input from more glomeruli. **e**, Comparing networks with or without PN-KC stereotypy. Performance was significantly worse when PN connections were correlated with one another. In the sparse condition, PN-KC connections were fixed to be sparse (K=7) and were randomly sampled from 50 PNs. In the sparse and correlated condition, the 50 PNs are subdivided into 3 evenly sized groups (17, 17, 16), and each KC only samples from PNs within a group.

**Figure S5.**
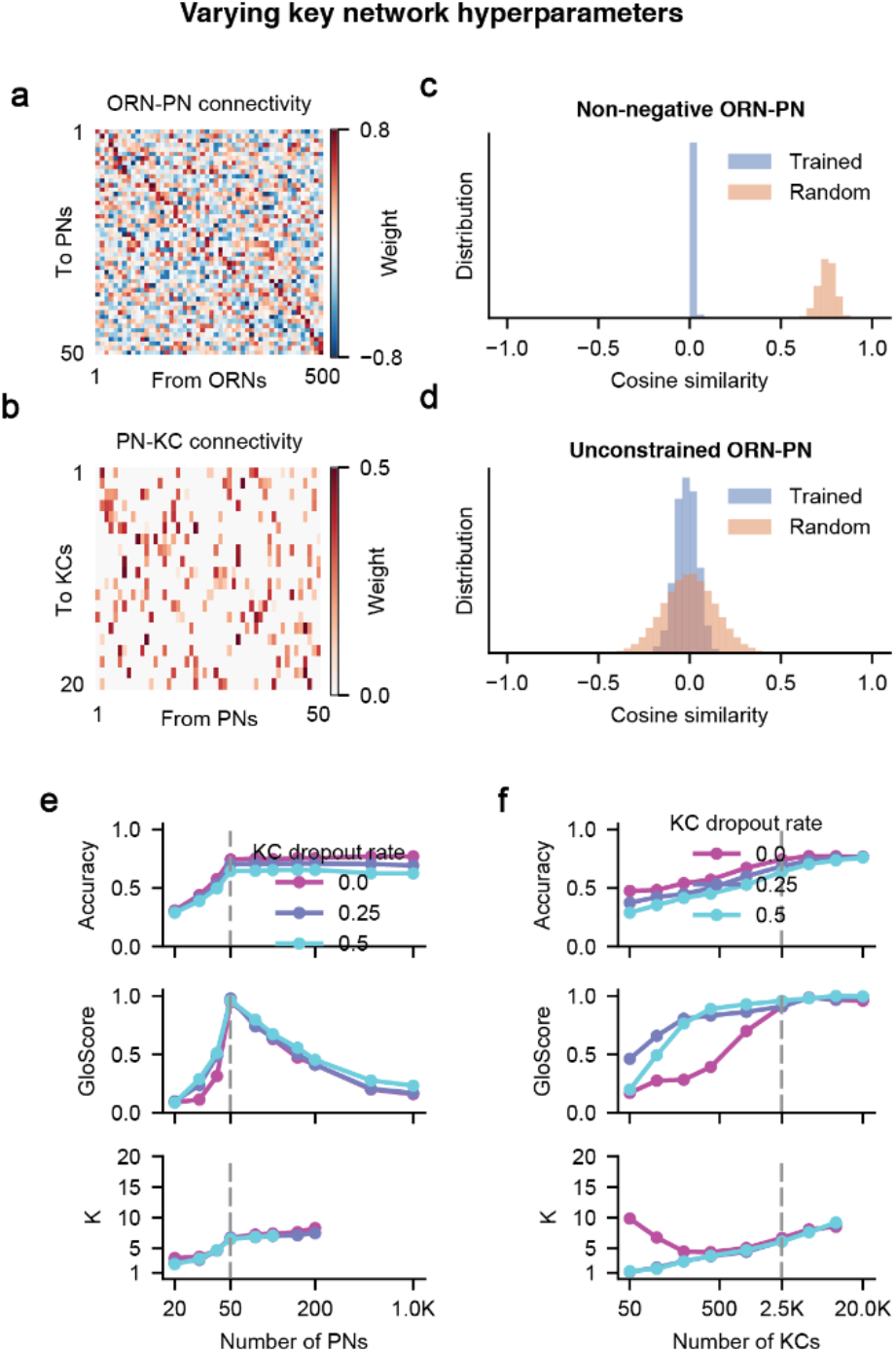
Impact of key network hyperparameters, Related to Figure 2. **a, b**, ORN-PN connectivity (**a**) and PN-KC connectivity (**b**) in a network without non-negative ORN-PN connections. **c, d**, Training decorrelates ORN-PN connections in networks with (**c**) and without (**d**) non-negative ORN-PN connections. Each PN unit receives connections from 500 ORNs, their weights summarized by a 500-dimensional vector. For two PN units, we compute the cosine similarity (cosine of angle) between each of their input weight vectors. The distribution of cosine similarities between all pairs of PN units in trained networks (blue), and in random networks (orange). If ORN-PN connections are non-negative, the random weights are drawn from a uniform distribution between 0 and 1, otherwise they are drawn from a random Gaussian distribution. These results show that training reduces the cosine similarity between input weights to pairs of PNs, decorrelating PNs. **e, f**, Accuracy, GloScore, KC input degree for networks with different numbers of PNs (**e**) and KCs (**f**), and for different levels of KC dropout rate.

**Figure S6.**
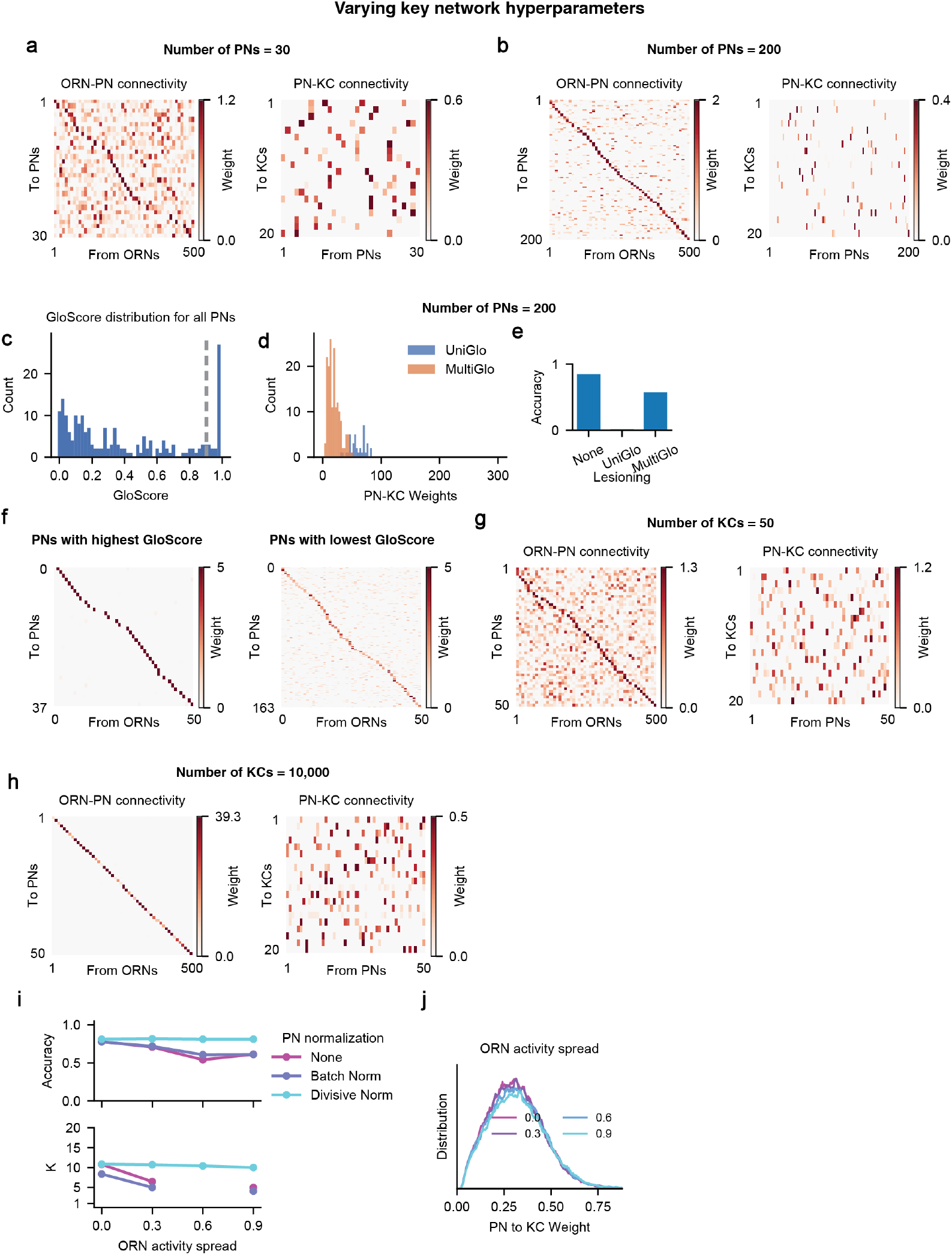
Impact of varying the number of neurons each layer and concentration datasets, Related to Figure 2. **a**, ORN-PN connectivity (left) and PN-KC connectivity (right) for a network with 30 PNs. **b**, Similar to (**a**), but for a network with 200 PNs. In neither case does ORN-PN connections form clean glomeruli. **c-f**, Analysis of a network with 200 PNs. **c**, The distribution of GloScore computed for each PN unit. A proportion of PNs have close to 1 GloScore. The threshold (dotted grey line) separates UniGlo units and MultiGlo units. **d**, UniGlo PN units tend to make stronger connections to KCs. **e**, Lesioning UniGlo PN units has a far stronger impact on classification accuracy. **f**, Connections from ORNs to PN with highest GloScore (left) and lowest GloScore (right). **g-h**, Similar to (**a, b**), but for networks with 50 and 10,000 KCs. **i**, Accuracy and KC input degree *K* for networks using different PN normalization and trained on different concentration datasets (see Figure 2d-e). Only PN-KC connections were trained in these networks. **j**, The distribution of PN-to-KC weights for networks using divisive normalization.

**Figure S7.**
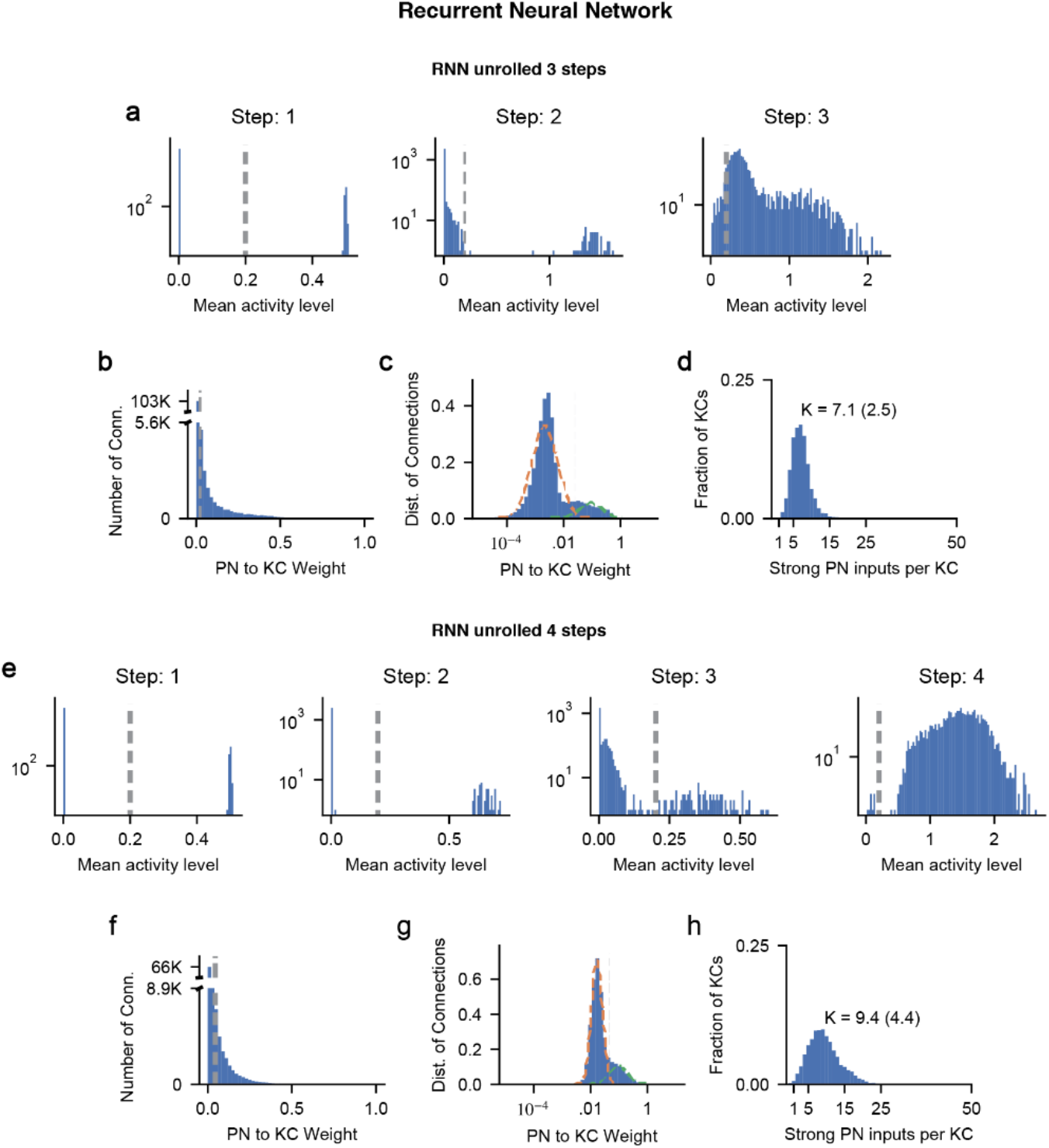
Additional analyses of the recurrent neural network model, Related to Figure 3. **a-d**. Analysis of a RNN unrolled for 3 steps. (**a**) Distribution of neuron’s mean activity level, computed at different processing steps. Left to right, step 1 to 3. For each neuron, we compute the average activity across all odors. Each dashed line corresponds to the threshold used to define active neurons. The same value of 0.2 used for all distributions. (**b,c**) Distribution of step 1 to step 2 (‘PN-KC’) connection weights after training, in linear space (**b**) and log space (**c**). No weight pruning is used for RNNs. In log space, the distribution is fit by a bi-modal Gaussian distribution. Strong PN-KC weights refer to the connections above the threshold separating these two modes. **d**, The distribution of strong “PN inputs to KC”. Here PN neurons refer to neurons active at step 1, while KC neurons refer to those active at step 2. **e-h**, Similar to (**a-d**), but for networks unrolled for 4 steps instead of 3. Classification readout occurs at step 4. Here “PN-KC weights” refers to the effective step 2-4 connectivity, which is the matrix product of the step 2-3 and step 3-4 connectivity.

**Figure S8.**
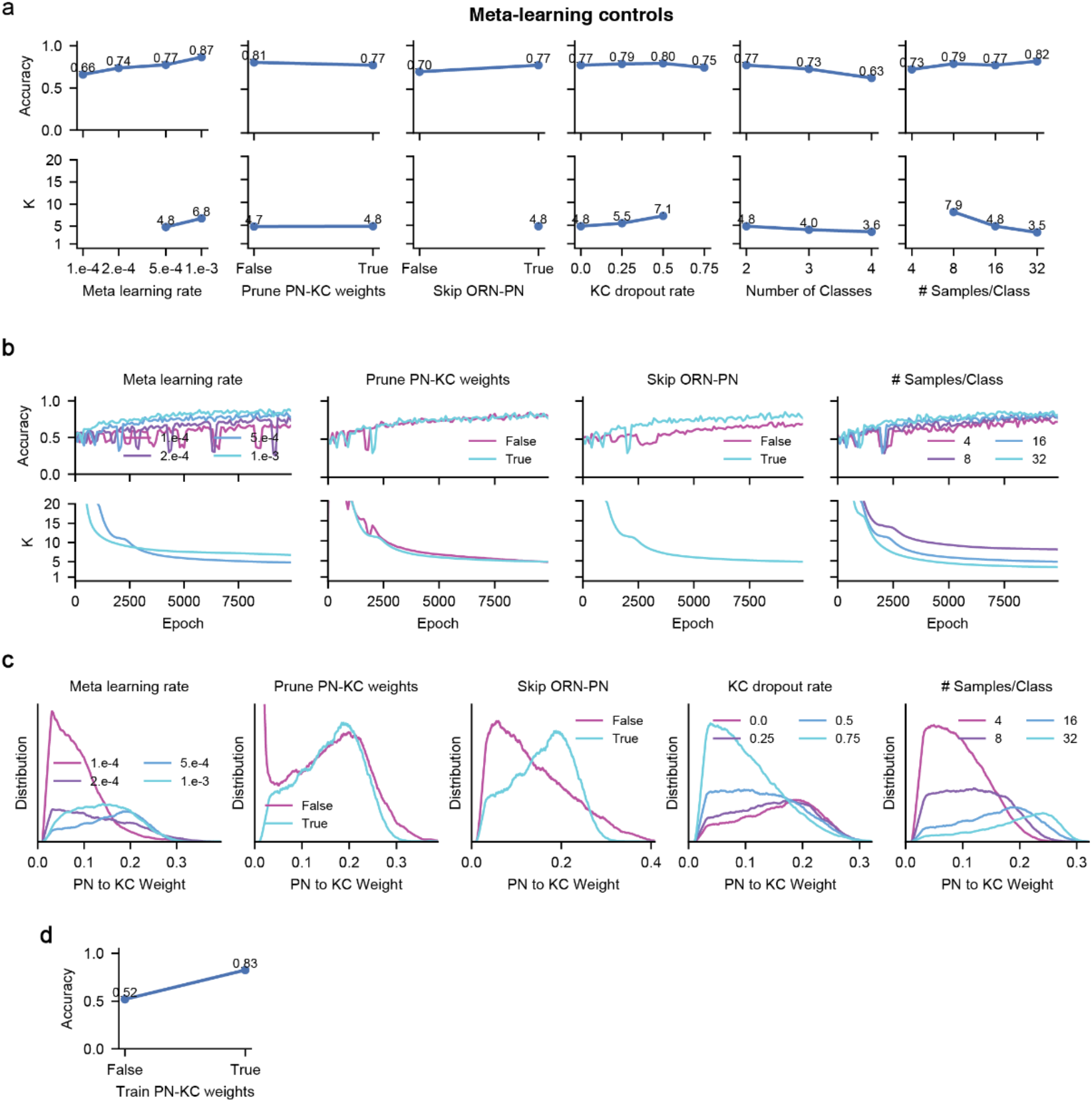
Formation of sparse connectivity during meta-learning, Related to Figure 4. **a**, Accuracy and KC input degree for networks meta-trained with different hyperparameter values. From left to right, meta-learning rate, whether weak PN-KC weights are pruned, whether to include a trainable ORN-PN layer, KC dropout rate, number of classes within each meta-learning episode (see Methods), number of samples per class within each episode. By default, the PN layer forms exact glomeruli (each PN unit receives connections only from the same type of ORNs), and the ORN-PN connections are fixed. Although PN-KC connectivity remains sparse, KC input degree is moderately affected by hyperparameter choices. Convention is the same as Figure S1. **b**, Accuracy and KC input degree during meta-training for networks with different hyperparameter values. **c**, Distribution of PN-KC weights after meta-training for different networks. **d**, Accuracy is around chance level (0.5) when PN-KC weights are fixed and not being meta-trained.

**Figure S9.**
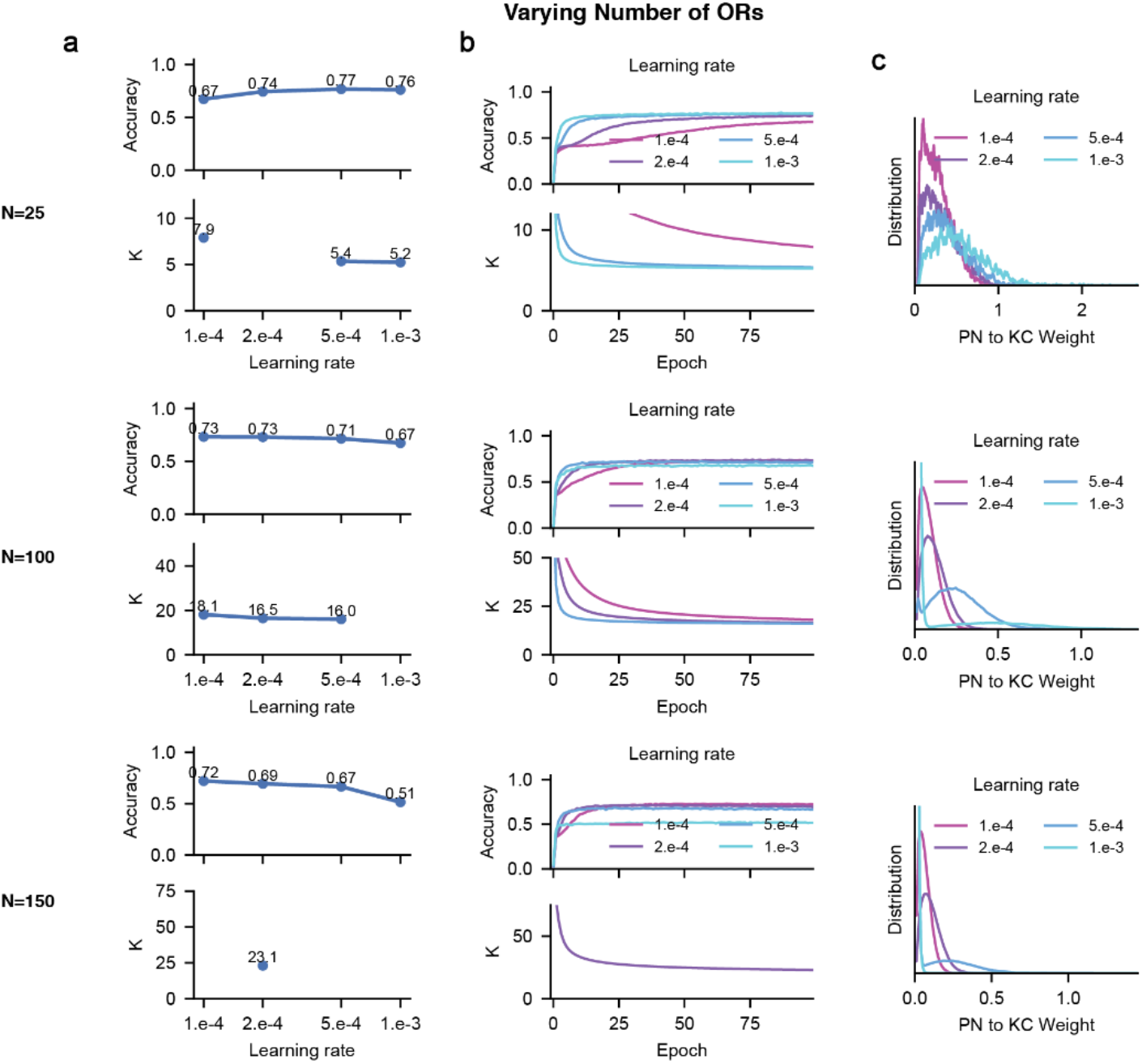
Impact of changing the number of olfactory receptors (ORs), Related to Figure 5. **a**, Accuracy and KC input degree for networks trained with different learning rates. From top to bottom, networks with different number of ORs (25, 100, 150). ORN-PN connectivity is fixed and PNs form exact glomeruli. Here we use PN to refer to the second compression layer in the network, and KC as the third expansion layer. **b**, Accuracy and KC input degree across training. **c**, The distributions of PN-KC weights for different learning rate values.

**Figure S10.**
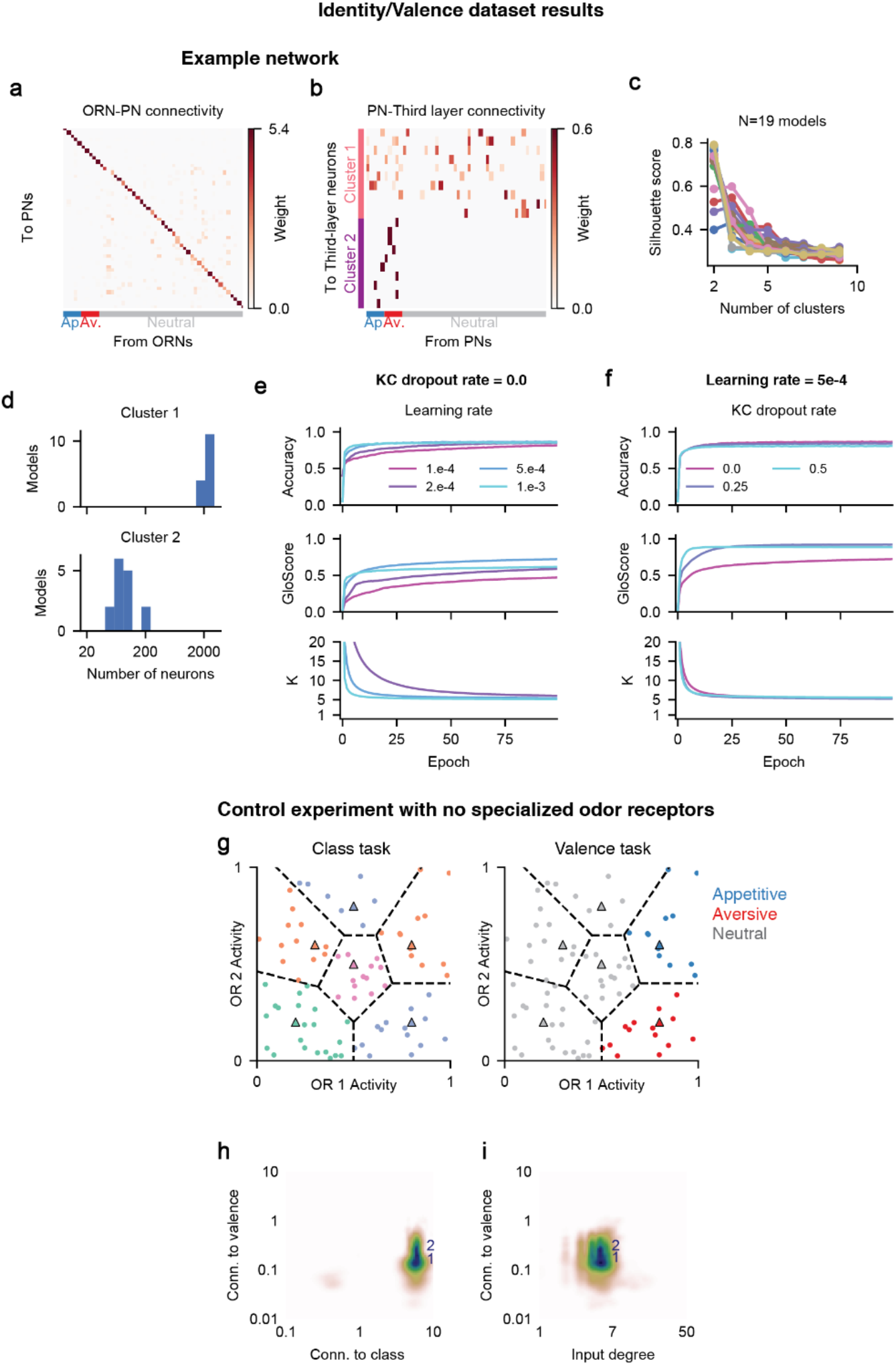
Additional analyses for the emergence of innate and learned pathways, Related to Figure 6. **a, b**, Connectivity of an example network. ORN-PN connectivity (**a**) and PN-Third layer connectivity (**b**). (**b**) is the same as Figure 3d. **c**, Silhouette score as a function of the number of clusters used for K-means clustering algorithm. Silhouette score rates how well the clusters segregate by comparing intra-cluster distance with inter-cluster distance. Across networks with different hyperparameter combinations (see Methods), Silhouette score peaks at number of cluster equals to 2 or, less commonly, 3. **d**, Across models, cluster 1 has around 2,000 neurons while cluster 2 has less than 200 neurons. Clusters are sorted according to their average connection strength to the valence classification head. So cluster 1 neurons has on average weaker connections to the valence output than cluster 2 neurons. Only networks in which the Silhouette score peaks at two clusters were analyzed (**c**). **e, f**, Accuracy, GloScore, KC input degree during training for different values of learning rates (**e**) and KC dropout rate (**f**). **g-i**, Results on a dataset with no specialized odor receptors. **g**, Innate odors no longer activate specialized receptors strongly. Compare with Figure 6a. **h**, **i**, Similar to Figure 6c, e, but for a network trained on the dataset with no specialized receptors. No clustering emerges, despite forcing the number of clusters to be 2 in the clustering algorithm.

## References

Abadi, M., Barham, P., Chen, J., Chen, Z., Davis, A., Dean, J., … Zheng, X. (2016). TensorFlow: A system for large-scale machine learning. Proceedings of the 12th USENIX Symposium on Operating Systems Design and Implementation, OSDI 2016, 265–283. Retrieved from https://arxiv.org/abs/1605.08695v2

Aso, Y., Hattori, D., Yu, Y., Johnston, R. M., Iyer, N. A., Ngo, T.-T., … Rubin, G. M. (2014a). The neuronal architecture of the mushroom body provides a logic for associative learning. ELife, 3. https://doi.org/10.7554/eLife.04577

Aso, Y., Hattori, D., Yu, Y., Johnston, R. M., Iyer, N. A., Ngo, T. T. B., … Rubin, G. M. (2014b). The neuronal architecture of the mushroom body provides a logic for associative learning. ELife. https://doi.org/10.7554/eLife.04577

Babadi, B., & Sompolinsky, H. (2014). Sparseness and Expansion in Sensory Representations. Neuron, 83(5), 1213–1226. https://doi.org/10.1016/j.neuron.2014.07.035

Bolding, K. A., & Franks, K. M. (2018). Recurrent cortical circuits implement concentration-invariant odor coding. Science, 361(6407). https://doi.org/10.1126/SCIENCE.AAT6904

Bottou, L. (2010). Large-scale machine learning with stochastic gradient descent. Proceedings of COMPSTAT 2010 - 19th International Conference on Computational Statistics, Keynote, Invited and Contributed Papers, 177–186. https://doi.org/10.1007/978-3-7908-2604-3_16

Buck, L., & Axel, R. (1991). A novel multigene family may encode odorant receptors: a molecular basis for odor recognition. Cell, 65(1), 175–187.

Cario, M. C., & Nelson, B. L. (1997). Modeling and generating random vectors with arbitrary marginal distributions and correlation matrix. Industrial Engineering.

Caron, S. J. C., Ruta, V., Abbott, L. F., & Axel, R. (2013). Random convergence of olfactory inputs in the Drosophila mushroom body. Nature. https://doi.org/10.1038/nature12063

Chia, J., & Scott, K. (2019). Activation of specific mushroom body output neurons inhibits proboscis extension and feeding behavior. BioRxiv. https://doi.org/10.1101/768960

Cohn, R., Morantte, I., & Ruta, V. (2015). Coordinated and Compartmentalized Neuromodulation Shapes Sensory Processing in Drosophila. Cell, 163(7), 1742–1755. https://doi.org/10.1016/j.cell.2015.11.019

Cueva, C. J., Wang, P. Y., Chin, M., & Wei, X. X. (2019). Emergence of functional and structural properties of the head direction system by optimization of recurrent neural networks. ArXiv.

Datta, S. R., Vasconcelos, M. L., Ruta, V., Luo, S., Wong, A., Demir, E., … Axel, R. (2008). The Drosophila pheromone cVA activates a sexually dimorphic neural circuit. Nature, 452(7186), 473–477. https://doi.org/10.1038/nature06808

Davison, I. G., & Ehlers, M. D. (2011). Neural circuit mechanisms for pattern detection and feature combination in olfactory cortex. Neuron, 70(1), 82–94.

Dayan, P., & Abbot, L. F. (2005). Theoretical Neuroscience: Computational and Mathematical Modeling of Neural Systems. Retrieved from https://mitpress.ublish.com/book/theoretical-neuroscience#tab-98

De Belle, J. S., & Heisenberg, M. (1994). Associative odor learning in Drosophila abolished by chemical ablation of mushroom bodies. Science. https://doi.org/10.1126/science.8303280

Dubnau, J., Grady, L., Kitamoto, T., & Tully, T. (2001). Disruption of neurotransmission in Drosophila mushroom body blocks retrieval but not acquisition of memory. Nature. https://doi.org/10.1038/35078077

Dweck, H. K. M., Ebrahim, S. A. M., Thoma, M., Mohamed, A. A. M., Keesey, I. W., Trona, F., … Hansson, B. S. (2015). Pheromones mediating copulation and attraction in Drosophila. Proceedings of the National Academy of Sciences of the United States of America. https://doi.org/10.1073/pnas.1504527112

Ebrahim, S. A. M., Dweck, H. K. M., Stökl, J., Hofferberth, J. E., Trona, F., Weniger, K., … Knaden, M. (2015). Drosophila Avoids Parasitoids by Sensing Their Semiochemicals via a Dedicated Olfactory Circuit. PLoS Biology. https://doi.org/10.1371/journal.pbio.1002318

Felsenberg, J., Jacob, P. F., Walker, T., Barnstedt, O., Edmondson-Stait, A. J., Pleijzier, M. W., … Waddell, S. (2018). Integration of Parallel Opposing Memories Underlies Memory Extinction. Cell. https://doi.org/10.1016/j.cell.2018.08.021

Finn, C., Abbeel, P., & Levine, S. (2017). Model-Agnostic Meta-Learning for Fast Adaptation of Deep Networks.

Franks, K. M., Russo, M. J., Sosulski, D. L., Mulligan, A., Siegelbaum, S. A., & Axel, R. (2011). Recurrent Circuitry Dynamically Shapes the Activation of Piriform Cortex. Neuron, 72(1), 49–56. https://doi.org/10.1016/j.neuron.2011.08.020

Godfrey, P. A., Malnic, B., & Buck, L. B. (2004). The mouse olfactory receptor gene family. Proceedings of the National Academy of Sciences, 101(7), 2156–2161. https://doi.org/10.1073/PNAS.0308051100

Gruntman, E., & Turner, G. C. (2013). Integration of the olfactory code across dendritic claws of single mushroom body neurons. Nature Neuroscience, 16(12), 1821–1829. https://doi.org/10.1038/nn.3547

Handler, A., Graham, T. G. W., Cohn, R., Morantte, I., Siliciano, A. F., Zeng, J., … Ruta, V. (2019). Distinct Dopamine Receptor Pathways Underlie the Temporal Sensitivity of Associative Learning. Cell. https://doi.org/10.1016/j.cell.2019.05.040

Hattori, D., Aso, Y., Swartz, K. J., Rubin, G. M., Abbott, L. F., & Axel, R. (2017). Representations of Novelty and Familiarity in a Mushroom Body Compartment. Cell. https://doi.org/10.1016/j.cell.2017.04.028

Heisenberg, M., Borst, A., Wagner, S., & Byers, D. (1985). drosophila mushroom body mutants are deficient in olfactory learning: Research papers. Journal of Neurogenetics. https://doi.org/10.3109/01677068509100140

Hige, T., Aso, Y., Rubin, G. M., & Turner, G. C. (2015). Plasticity-driven individualization of olfactory coding in mushroom body output neurons. Nature, 526(7572), 258–262. https://doi.org/10.1038/nature15396

Ioffe, S., & Szegedy, C. (2015). Batch normalization: Accelerating deep network training by reducing internal covariate shift. 32nd International Conference on Machine Learning, ICML 2015.

Jefferis, G. S. X. E., Potter, C. J., Chan, A. M., Marin, E. C., Rohlfing, T., Maurer, C. R., & Luo, L. (2007). Comprehensive Maps of Drosophila Higher Olfactory Centers: Spatially Segregated Fruit and Pheromone Representation. Cell. https://doi.org/10.1016/j.cell.2007.01.040

Kazama, H., & Wilson, R. I. (2009). Origins of correlated activity in an olfactory circuit. Nature Neuroscience, 12(9), 1136–1144. https://doi.org/10.1038/nn.2376

Kingma, D. P., & Ba, J. (2014). Adam: A Method for Stochastic Optimization. Retrieved from http://arxiv.org/abs/1412.6980

Kurtovic, A., Widmer, A., & Dickson, B. J. (2007). A single class of olfactory neurons mediates behavioural responses to a Drosophila sex pheromone. Nature. https://doi.org/10.1038/nature05672

Lecun, Y., Bengio, Y., & Hinton, G. (2015). Deep learning. Nature. https://doi.org/10.1038/nature14539

Li, F., Lindsey, J., Marin, E. C., Otto, N., Dreher, M., Dempsey, G., … Rubin, G. M. (2020). The connectome of the adult drosophila mushroom body provides insights into function. ELife. https://doi.org/10.7554/eLife.62576

Lin, A. C., Bygrave, A. M., De Calignon, A., Lee, T., & Miesenböck, G. (2014). Sparse, decorrelated odor coding in the mushroom body enhances learned odor discrimination. Nature Neuroscience, 17(4), 559–568. https://doi.org/10.1038/nn.3660

Litwin-Kumar, A., Harris, K. D., Axel, R., Sompolinsky, H., & Abbott, L. F. (2017). Optimal Degrees of Synaptic Connectivity. Neuron, 93(5), 1153–1164.e7. https://doi.org/10.1016/j.neuron.2017.01.030

Luo, S. X., Axel, R., & Abbott, L. F. (2010). Generating sparse and selective third-order responses in the olfactory system of the fly. Proceedings of the National Academy of Sciences, 107(23), 10713–10718. https://doi.org/10.1073/pnas.1005635107

Mante, V., Sussillo, D., Shenoy, K. V., & Newsome, W. T. (2013). Context-dependent computation by recurrent dynamics in prefrontal cortex. Nature. https://doi.org/10.1038/nature12742

Marin, E. C., Jefferis, G. S. X. E., Komiyama, T., Zhu, H., & Luo, L. (2002). Representation of the glomerular olfactory map in the Drosophila brain. Cell. https://doi.org/10.1016/S0092-8674(02)00700-6

Marr, D. (1969). A theory of cerebellar cortex. The Journal of Physiology, 202(2), 437–470. Retrieved from http://www.ncbi.nlm.nih.gov/pubmed/5784296

Masse, N. Y., Yang, G. R., Song, H. F., Wang, X. J., & Freedman, D. J. (2019). Circuit mechanisms for the maintenance and manipulation of information in working memory. Nature Neuroscience. https://doi.org/10.1038/s41593-019-0414-3

Masuda-Nakagawa, L. M., Tanaka, N. K., & O’Kane, C. J. (2005). Stereotypic and random patterns of connectivity in the larval mushroom body calyx of Drosophila. Proceedings of the National Academy of Sciences of the United States of America. https://doi.org/10.1073/pnas.0509643102

McGuire, S. E., Le, P. T., & Davis, R. L. (2001). The role of Drosophila mushroom body signaling in olfactory memory. Science. https://doi.org/10.1126/science.1062622

Min, S., Ai, M., Shin, S. A., & Suh, G. S. B. (2013). Dedicated olfactory neurons mediating attraction behavior to ammonia and amines in Drosophila. Proceedings of the National Academy of Sciences of the United States of America. https://doi.org/10.1073/pnas.1215680110

Miyamichi, K., Amat, F., Moussavi, F., Wang, C., Wickersham, I., Wall, N. R., … Luo, L. (2011). Cortical representations of olfactory input by trans-synaptic tracing. Nature, 472(7342), 191–196. https://doi.org/10.1038/nature09714

Mombaerts, P., Wang, F., Dulac, C., Chao, S. K., Nemes, A., Mendelsohn, M., … Axel, R. (1996). Visualizing an Olfactory Sensory Map. Cell, 87(4), 675–686. https://doi.org/10.1016/S0092-8674(00)81387-2

Oliphant, T. E. (2006). Guide to NumPy. Retrieved from http://www.trelgol.com

Olsen, S. R., Bhandawat, V., & Wilson, R. I. (2010). Divisive normalization in olfactory population codes. Neuron, 66(2), 287–299. https://doi.org/10.1016/j.neuron.2010.04.009

Pashkovski, S. L., Iurilli, G., Brann, D., Chicharro, D., Drummey, K., Franks, K. M., … Datta, S. R. (2020). Structure and flexibility in cortical representations of odour space. Nature 2020 583:7815, 583(7815), 253–258. https://doi.org/10.1038/s41586-020-2451-1

Paszke, A., Gross, S., Massa, F., Lerer, A., Bradbury, J., Chanan, G., … Chintala, S. (2019). PyTorch: An Imperative Style, High-Performance Deep Learning Library. Advances in Neural Information Processing Systems, 32. Retrieved from https://arxiv.org/abs/1912.01703v1

Pedregosa, F., Varoquaux, G., Gramfort, A., Michel, V., Thirion, B., Grisel, O., … Duchesnay, É. (2011). Scikit-learn: Machine Learning in Python. Journal of Machine Learning Research, 12(85), 2825–2830. Retrieved from http://jmlr.org/papers/v12/pedregosa11a.html

Price, J. L., & Powell, T. P. (1970). The mitral and short axon cells of the olfactory bulb. J Cell Sci, 7(3), 631–651. Retrieved from https://www.ncbi.nlm.nih.gov/pubmed/5492279

Reardon, T. R., Murray, A. J., Turi, G. F., Wirblich, C., Croce, K. R., Schnell, M. J., … Losonczy, A. (2016). Rabies Virus CVS-N2cδG Strain Enhances Retrograde Synaptic Transfer and Neuronal Viability. Neuron. https://doi.org/10.1016/j.neuron.2016.01.004

Ressler, K. J., Sullivan, S. L., & Buck, L. B. (1993). A zonal organization of odorant receptor gene expression in the olfactory epithelium. Cell, 73(3), 597–609. https://doi.org/10.1016/0092-8674(93)90145-G

Ressler, K. J., Sullivan, S. L., & Buck, L. B. (1994). Information coding in the olfactory system: Evidence for a stereotyped and highly organized epitope map in the olfactory bulb. Cell, 79(7), 1245–1255. https://doi.org/10.1016/0092-8674(94)90015-9

Rumelhart, D. E., Hinton, G. E., & Williams, R. J. (1986). Learning representations by back-propagating errors. Nature, 323(6088), 533–536. https://doi.org/10.1038/323533a0

Ruta, V., Datta, S. R., Vasconcelos, M. L., Freeland, J., Looger, L. L., & Axel, R. (2010). A dimorphic pheromone circuit in Drosophila from sensory input to descending output. Nature. https://doi.org/10.1038/nature09554

Scheffer, L. K., Xu, C. S., Januszewski, M., Lu, Z., Takemura, S. Y., Hayworth, K. J., … Plaza, S. M. (2020). A connectome and analysis of the adult drosophila central brain. ELife. https://doi.org/10.7554/ELIFE.57443

Schoonover, C. E., Ohashi, S. N., Axel, R., & Fink, A. J. P. (2021). Representational drift in primary olfactory cortex. Nature 2021 594:7864, 594(7864), 541–546. https://doi.org/10.1038/s41586-021-03628-7

Stensmyr, M. C., Dweck, H. K. M., Farhan, A., Ibba, I., Strutz, A., Mukunda, L., … Hansson, B. S. (2012). A conserved dedicated olfactory circuit for detecting harmful microbes in drosophila. Cell. https://doi.org/10.1016/j.cell.2012.09.046

Stern, M., Bolding, K. A., Abbott, L. F., & Franks, K. M. (2018). A transformation from temporal to ensemble coding in a model of piriform cortex. ELife, 7. https://doi.org/10.7554/ELIFE.34831

Suh, G. S. B., Wong, A. M., Hergarden, A. C., Wang, J. W., Simon, A. F., Benzer, S., … Anderson, D. J. (2004). A single population of olfactory sensory neurons mediates an innate avoidance behaviour in Drosophila. Nature. https://doi.org/10.1038/nature02980

Tanaka, N. K., Awasaki, T., Shimada, T., & Ito, K. (2004). Integration of chemosensory pathways in the Drosophila second-order olfactory centers. Current Biology. https://doi.org/10.1016/j.cub.2004.03.006

Tanaka, N. K., Tanimoto, H., & Ito, K. (2008). Neuronal assemblies of the Drosophila mushroom body. Journal of Comparative Neurology. https://doi.org/10.1002/cne.21692

Uria, B., Ibarz, B., Banino, A., Zambaldi, V., Kumaran, D., Hassabis, D., … Blundell, C. (2020). The spatial memory pipeline: A model of egocentric to allocentric understanding in mammalian brains. BioRxiv. https://doi.org/10.1101/2020.11.11.378141

Varela, N., Gaspar, M., Dias, S., & Vasconcelos, M. L. (2019). Avoidance response to CO 2 in the lateral horn. PLoS Biology. https://doi.org/10.1371/journal.pbio.2006749

Vassar, R., Chao, S. K., Sitcheran, R., Nuñez, J. M., Vosshall, L. B., & Axel, R. (1994). Topographic organization of sensory projections to the olfactory bulb. Cell, 79(6), 981–991.

Virtanen, P., Gommers, R., Oliphant, T. E., Haberland, M., Reddy, T., Cournapeau, D., … van Mulbregt, P. (2020). SciPy 1.0: fundamental algorithms for scientific computing in Python. Nature Methods 2020 17:3, 17(3), 261–272. https://doi.org/10.1038/s41592-019-0686-2

Vosshall, L. B., Amrein, H., Morozov, P. S., Rzhetsky, A., & Axel, R. (1999). A spatial map of olfactory receptor expression in the Drosophila antenna. Cell. https://doi.org/10.1016/S0092-8674(00)80582-6

Vosshall, L. B., Wong, A. M., & Axel, R. (2000a). An olfactory sensory map in the fly brain. Cell. https://doi.org/10.1016/S0092-8674(00)00021-0

Vosshall, L. B., Wong, A. M., & Axel, R. (2000b). An Olfactory Sensory Map in the Fly Brain. Cell, 102(2), 147–159. https://doi.org/10.1016/S0092-8674(00)00021-0

Wilson, R. I. (2013). Early Olfactory Processing in Drosophila : Mechanisms and Principles. Annual Review of Neuroscience. https://doi.org/10.1146/annurev-neuro-062111-150533

Wong, A. M., Wang, J. W., & Axel, R. (2002). Spatial representation of the glomerular map in the Drosophila protocerebrum. Cell. https://doi.org/10.1016/S0092-8674(02)00707-9

Yamins, D. L. K., & DiCarlo, J. J. (2016). Using goal-driven deep learning models to understand sensory cortex. Nature Neuroscience. https://doi.org/10.1038/nn.4244

Yamins, D. L. K., Hong, H., Cadieu, C. F., Solomon, E. A., Seibert, D., & DiCarlo, J. J. (2014). Performance-optimized hierarchical models predict neural responses in higher visual cortex. Proceedings of the National Academy of Sciences of the United States of America. https://doi.org/10.1073/pnas.1403112111

Yang, G. R., Joglekar, M. R., Song, H. F., Newsome, W. T., & Wang, X. J. (2019). Task representations in neural networks trained to perform many cognitive tasks. Nature Neuroscience. https://doi.org/10.1038/s41593-018-0310-2

Zhang, X., & Firestein, S. (2002). The olfactory receptor gene superfamily of the mouse. Nat Neurosci, 5(2), 124–133. https://doi.org/10.1038/nn800

Zheng, Z., Lauritzen, J. S., Perlman, E., Robinson, C. G., Nichols, M., Milkie, D., … Bock, D. D. (2018). A Complete Electron Microscopy Volume of the Brain of Adult Drosophila melanogaster. Cell. https://doi.org/10.1016/j.cell.2018.06.019

