## Supplementary material for "Evolving the olfactory system with machine learning": Video_S1_legend

**Convergence of like ORNs onto unique PNs and emergence of sparse PN-KC connectivity during network training**

Visualization of ORN-PN and PN-KC connectivity during network training. The opacity of each line segment is proportional to the connection weight. Only a subset of the network is shown for clarity. ORNs that express the first 3 of 50 unique ORs are shown in the ORN layer. All 50 PNs are shown in the PN layer. The first 5 of 2500 KCs are shown in the KC layer. ORN-PN connections are colored according to the identity of the OR. PN-KC connections are colored according to the identity of the KC.
